# Identifying potential natural inhibitors of *Brucella melitensis* Methionyl-tRNA synthetase through an *in-silico* approach

**DOI:** 10.1101/2021.09.09.459562

**Authors:** Adekunle Babjide Rowaiye, Akwoba Joseph Ogugua, Gordon Ibeanu, Doofan Bur, Osaretin Benjamin Ogbeide, Emmanuella Oshiorenimeh Abraham, Hamzah Bundu Usman

## Abstract

**Background:** Brucellosis is an infectious disease caused by bacteria of the genus *Brucella*. Although it is the most common zoonosis worldwide, there are increasing reports of drug resistance and cases of relapse after long term treatment with the existing drugs of choice. This study therefore aims at identifying possible natural inhibitors of *Brucella melitensis* Methionyl-tRNA synthetase through an *in-silico* approach.

**Methods:** Using PyRx 0.8 virtual screening software, the target was docked against a library of natural compounds obtained from edible African plants. The compound, 2-({3-[(3,5-dichlorobenzyl) amino] propyl} amino) quinolin-4(1H)-one (OOU) which is a co-crystallized ligand with the target was used as the reference compound. Screening of the molecular descriptors of the compounds for bioavailability, pharmacokinetic properties, and bioactivity was performed using the SWISSADME, pkCSM, and Molinspiration web servers respectively. The Fpocket and PLIP webservers were used to perform the analyses of the binding pockets and the protein ligand interactions. Analysis of the time-resolved trajectories of the Apo and Holo forms of the target was performed using the Galaxy and MDWeb servers. The lead compounds, Strophanthidin and Isopteropodin are present in *Corchorus olitorius* and *Uncaria tomentosa* (cat-claw) plants respectively.

**Results:** Isopteropodin had a binding affinity score of -8.9 kcal / ml with the target and had 17 anti-correlating residues in pocket 1 after molecular dynamics simulation. The complex formed by Isopteropodin and the target had a total RMSD of 4.408 and a total RMSF of 9.8067. However, Strophanthidin formed 3 hydrogen bonds with the target at ILE21, GLY262 and LEU294, and induced a total RMSF of 5.4541 at Pocket 1.

**Conclusion:** Overall, Isopteropodin and Strophanthidin were found to be better drug candidates than OOU and they showed potentials to inhibit the *Brucella melitensis* Methionyl-tRNA synthetase at Pocket 1, hence abilities to treat brucellosis. *In vivo* and *in vitro* investigations are needed to further evaluate the efficacy and toxicity of the lead compounds.

**Author Summary:**

1. Strophanthidin and Isopteropodin showed potentials to inhibit the *Brucella melitensis* Methionyl-tRNA synthetase at Pocket 1
2. Both compounds can be used to treat brucellosis.
3. Both compounds showed potentials of being safe to use in humans.

## Introduction

Brucellosis is an infectious disease caused by bacteria of the genus *Brucella*. The species are Gram-negative intracellular coccobacilli that occur in a wide variety of animals including cattle, sheep, goats, pigs, other livestock as well as humans [1]. There are 12 species of *Brucella* based on specificity of host [2]. Although, *Brucella* species are often associated with certain hosts, they infect others apart from their preferred hosts. Being basically a disease of animals, most human brucellosis cases are traceable to infected animals or their products [3]. Hence, its control in human populations is targeted at the animals. Most infections in humans are due to contact with contaminated materials. The disease therefore has a major occupational disposition among livestock workers, veterinarians, abattoir workers, so also hides, skin and wool workers as well as laboratory personnel [4,5]. To the general public, brucellosis is mainly transmitted through the consumption of unpasteurized contaminated milk or its products [6,7]. In few occasions, human-to-human transmissions have been recorded through sexual contact, blood transfusion, bone marrow transplant, obstetrical manipulations during child birth and congenital means [8,4,9]. Brucellosis however is noted as the most common zoonosis worldwide with more than 500,000 cases recorded annually [10].

The disease is well controlled in most developed countries [11], but common in Africa, South America, Asia, the Caribbean, Middle East and the Mediterranean basin [12,2,13]. In livestock production, the major economic effects are due to abortion, premature birth, reduced milk production, repeat breeding and cost of veterinary care [14]. In humans, the disease results in loss of manpower as well as huge costs in medical care [15]. Thus, the control of the disease in most developed countries has resulted in significant economic gains as well as reduction in human cases. However, in developing countries the disease is still of major economic and public health importance. This is mainly due to lack of well-defined control policies as well as the lifestyle of high-risk persons who are mostly uninformed about the disease [16]. In controlling brucellosis, many countries embark on or consider the actions compatible with their tradition and resources. Methods of controlling brucellosis therefore are hinged on diagnosis, control, increasing the awareness of the disease and vaccination [17]. Brucellosis remains a largely neglected disease especially in developing countries [3]. In sub-Sahara Africa, there has been little attention towards the control and prevention of brucellosis except in South Africa [18]. The control of brucellosis in Africa is hindered by many factors. The farming system is basically traditional. Nomadism which accounts for as high as 95% of cattle production in many West African countries [3] involves uncontrolled movement of livestock: a major risk factor in the spread of brucellosis [16]. Brucellosis is therefore noted to impact negatively on human and animal health, hampers social and economic progress as well as food security in developing economies [19].

*Brucella* remains a potential bio-terroristic agent and moreover, treatment of the disease is quite difficult in affected people because of the ability of the organism to evade the host immune system and reside in the cell for extended periods [20]. Most drugs currently used to treat *Brucella* infection have not been relatively effective. This is because *Brucella* activates the cAMP/protein kinase A pathway which is crucial for the survival and establishment of *Brucella* within macrophages. Inside the cells, they inhibit programmed cell death leading to long survival in the cells. Also, prevalence of drug resistance genes is being reported in *Brucella* species [21,22]. Such reports of antibiotic resistance are rendering the use of antibiotics almost useless in treatment of brucellosis [23]. Therefore, cases of relapse after a long period of treatment are common [8]. This underscores the need to search for alternatives to the current long term combination chemotherapy of brucellosis with the drugs of choice, which are rifampin, streptomycin and doxycycline [24]. Such new agents need to be able to penetrate and function within the macrophage cytoplasm, be inexpensive, non-toxic and more effective than the drugs traditionally used to treat the disease.

Availability of sequenced genome of *Brucella* species has offered new options in the search for drugs targeting enzymes that could be of use due to pathogen-host physiological and biochemical differences. The methionyl-tRNA synthetase, which is a member of the aminoacyl tRNA synthetase, has been identified as being very important for its roles in protein synthesis due to its recognition of initiator tRNA and tRNA delivering methionine for protein chain elongation [25]. According to Ojo *et al*. [26], methionyl-tRNA synthetase is promising as a good target for brucellosis drug development. Therefore, lead compounds targeting the enzyme could be useful and offer good alternatives for the treatment of brucellosis. The aim of this study is to use in-silico method to identify compounds of plant origin that can inhibit the activity of *Brucella melitensis* methionyl-tRNA synthesis and serve as remedies for brucellosis.

## Method

The *SWISS-MODEL* homology modeling server was used to model the target protein after the crystal structure of methionyl-tRNA synthetase MetRS from *Brucella melitensis* (PDB ID: 5K0S.1.A) [27,28]. The structure of the target protein was visualized using PyMOL [29], analyzed using the VADAR 1.8 server [30] and validated using the MolProbity server [31].

A library of 1,524 phytoconstituents belonging to different classes of secondary metabolites was collected from the results of the phytochemical analyses of edible African plants found in literature. The Structure Data File (sdf) formats of the 3D chemical structures of these compounds were downloaded from the PubChem database [32]. All ligands were loaded on the PyRx 0.8 software, their geometries were minimized and they were converted into the pdbqt format in readiness for molecular docking [33].

Docking of all the ligands against the target was performed using the AutoDock Vina tool of PyRx 0.8 software. The grid parameters were set at Center - x: 26.4524, y: 19.1969, z: 22.6112 and Dimensions – x: 64.4178, y: 72.5182, z: 84.2900. The setting for the docking was the universal force field (*UFF*) and conjugate gradient algorithm [33]. The 2-({3-[(3,5-dichlorobenzyl) amino] propyl} amino) 6uinoline-4(1H)-one (OOU) (PubChem ID 18353708) which is the co-crystallized ligand of the target protein was used as the reference compound. From the docking results, all docking scores higher than the binding affinity score of OOU (reference compound) with the target were screened out.

The predictions for molar refractivity, saturation and promiscuity for the front runner compounds were obtained from the SwissADME server and screening was performed based on established medicinal chemistry criteria [34]. Screening for absorption, distribution, metabolism, elimination, and toxicity (ADMET) properties was performed using the pkCSM server [35]. Further screening of the front runner compounds for bioactivity was performed with the Molinspiration server [36]. The PLIP webserver was used to decipher the hydrogen bonds, halogen bonds and hydrophobic interactions between residues of the target and the lead compounds [37].

A molecular dynamic simulation study of the apo and holo forms of the target protein was performed using the Galaxy and MDWeb servers [38,39]. Analyses of the time-resolved trajectory were done using parameters such as root-mean-square deviation (RMSD), root mean square fluctuation (RMSF), radius of gyration (RoG), B-factor, principal component analysis (PCA), and dynamical cross-correlation matrix (DCCM) [38,39].

The fasta format of the amino acid sequences of the target protein (P59078) was obtained from the UniProtKB database [40]. Sequences were placed on the BLAST tool of the NCBI server and the settings were PDB protein for database, Homo sapiens (taxid 96906) for organism, and blastp for algorithm [41].

## Results

### Analysis of the structure of the target

The modelled target protein had 507 residues with a 100% similarity identity with *Brucella melitensis* methionyl-tRNA synthetase (BrMelMetRS) (PDB: 5K0S) and also a qualitative model energy analysis (QMEAN) value of 0.76 and global model quality estimate (GMQE) value of 0.96. Resolved by X-ray diffraction method, the crystal structure of BrMelMetRS (PDB: 5K0S) showed a resolution of 2.45 Å and R-Value Free of 0.256 (Figure 1). The secondary structures of the target included 49% alpha helix, 22% beta sheets and 28% coils. The total solvent-accessible surface area (SASA) was 22269.0 (Å)^2^. Ramanchandran analysis revealed that in terms of geometry, the target protein had 1.21% poor rotamers, 97.1% favoured rotamers of which 99.6% were in allowed regions, 0.40% Ramachandran outliers, 98.02% ramanchandran favoured, 0.00% Cβ deviations (>0.25Å), and Rama distribution Z-score of 1.11 ± 0.36, 0.07% bad bonds and 0.48% band angles (Figure 2). With regards to low-resolution criteria, there were 0.8% carbon-alpha based low-resolution annotation method (CaBLAM) outliers and 0.60% carbon-alpha geometry outliers.

**Figure 1:**
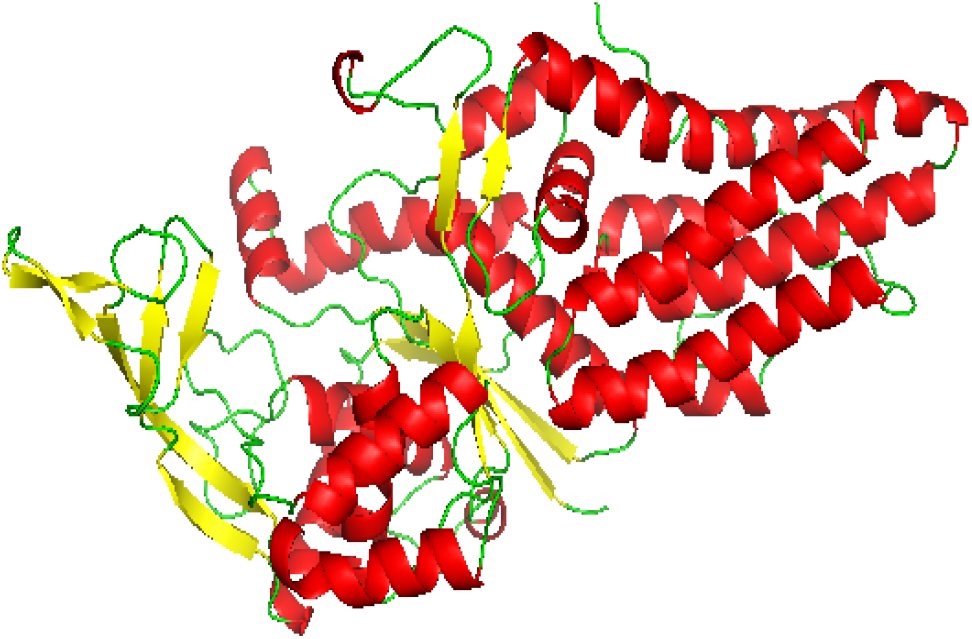
The cartoon structure of modeled BrMelMetRS. Beta sheets in yellow, alpha helix in red, and loops in green.

**Figure 2:**
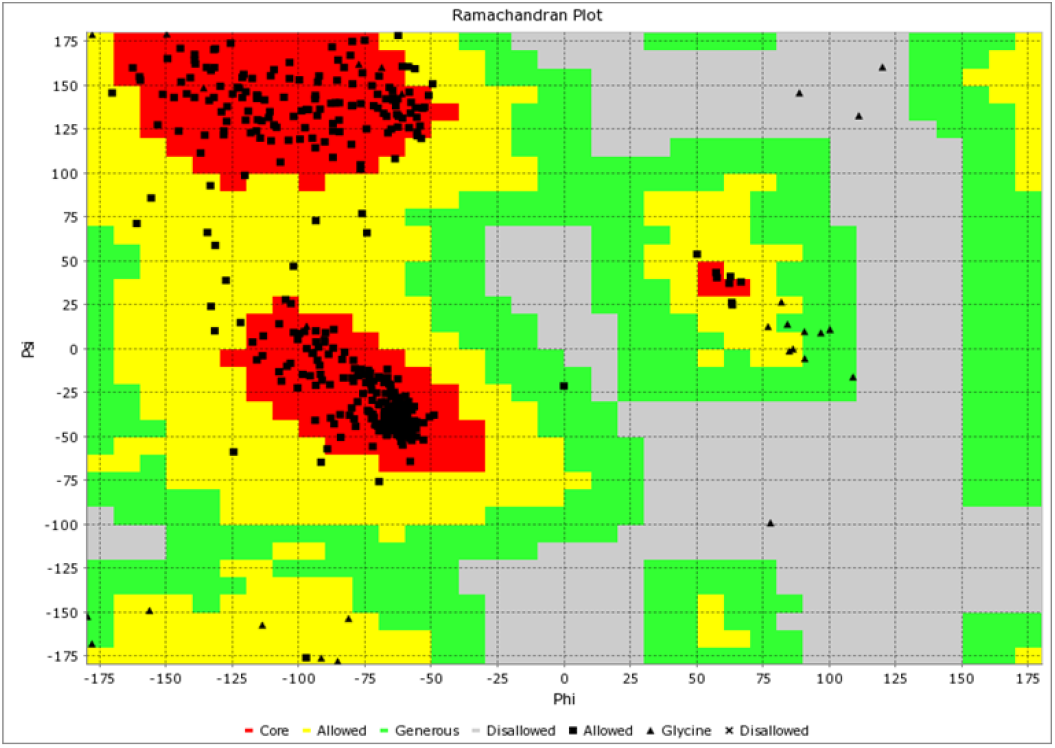
Ramachandran plot of modeled BrMelMetRS

### Drug-likeness properties and other molecular descriptors of ligands

For the reference and lead compounds, the drug-likeness properties such as hydrogen bond acceptor (HBA), hydrogen bond donor (HBD), log P, molecular weight, and topological surface area (TPSA) did not exceed 10, 5, 500 g/mol and 140 Å respectively (Figure 3 and Table 4). Furthermore, the reference and lead compounds’ molar refractivity ranged from 40 to 130, despite the fact that their number of rotatable bonds did not surpass 10. In terms of bioactivity, OOU and Strophanthidin had enzyme inhibition prediction values larger than 0.00, whereas Isopteropodin had a value less than 0.00. All the compounds had all their bioactivity prediction values greater than -5.00. From the bioavailability radars, all the compounds were within the range for drug-likeness properties of size, lipophilicity, solubility, polarity and flexibility (Figure 4). While OOU was slightly unsaturated, Strophanthidin and Isopteropodin were within the saturation range (above 0.25).

**Figure 3:**
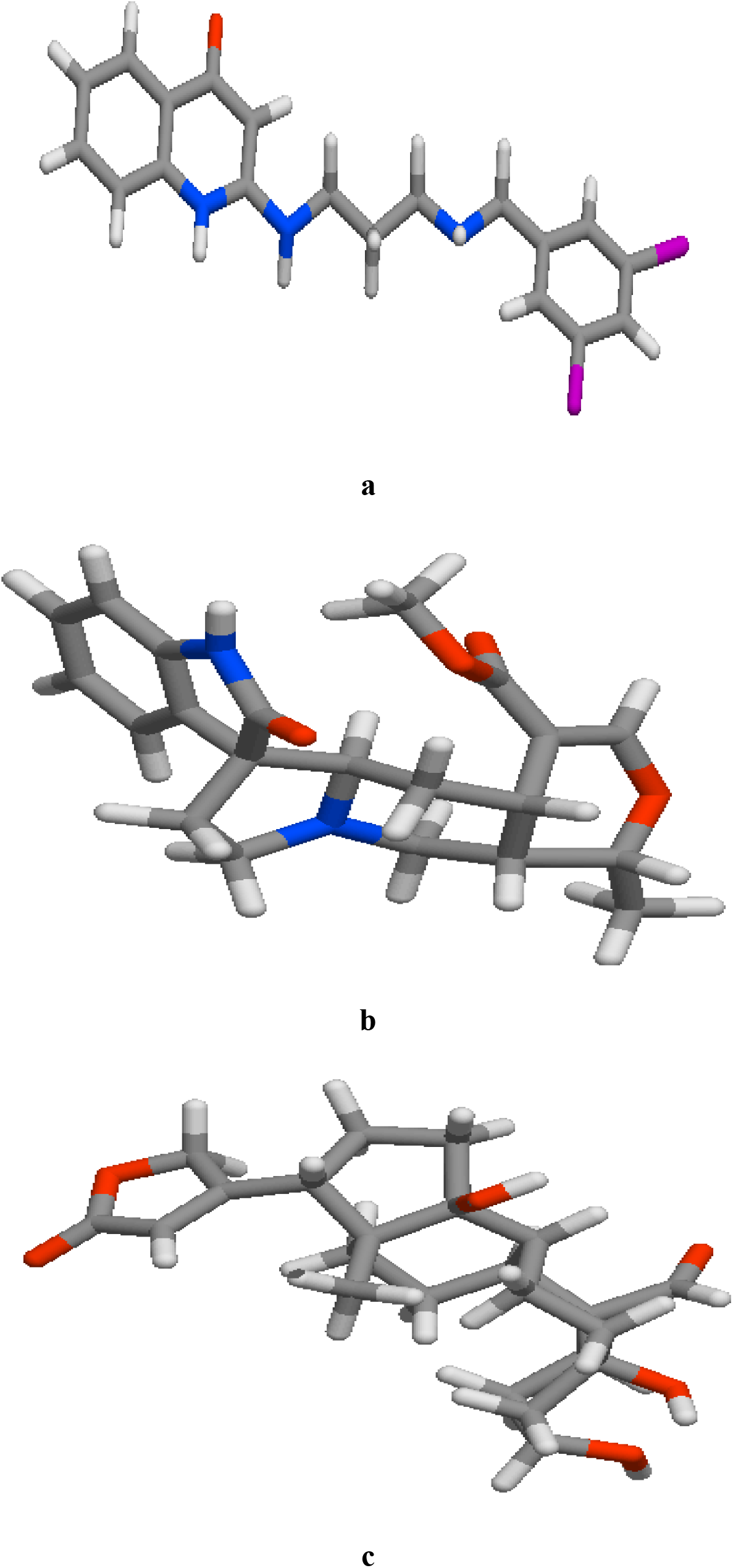
The stick model of the 3D structures of the reference and lead compounds (a) OOU (b) Isopteropodin (c) Strophanthidin

**Figure 4:**
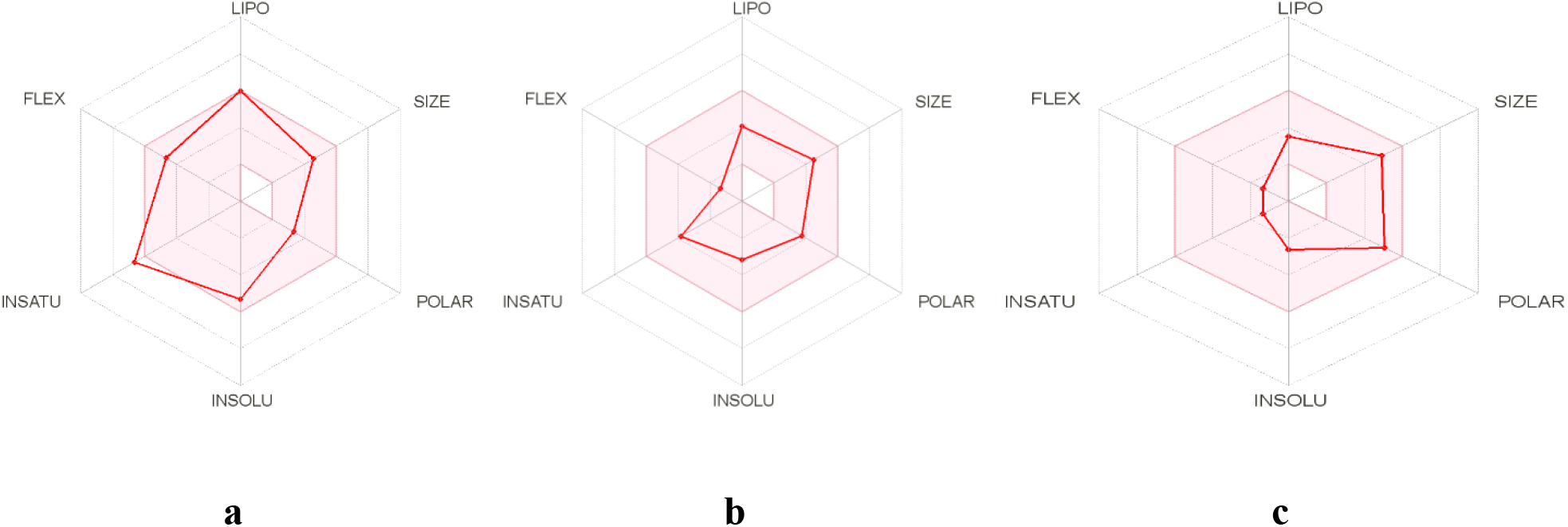
The bioavailability radars for reference and lead compounds (a) OOU (b) Isopteropodin (c) Strophanthidin

### The ADMET properties of ligands

From Table 2, the water solubility values for both the leads and reference compounds were greater than -6.0 log mol/L. The values for OOU and Isopteropodin’s caco2 permeability (log Papp in 10-6 cm.s-1) were larger than 0.9, while Strophanthidin‘s value was less than 0.9. For all of the compounds, the human intestine absorption (percentage absorbed) values were greater than 30%. Similarly, all the compounds had skin permeability (LogKp) values less than -2.5 (Table 2). Remarkably, OOU was predicted to be inhibitor of both P-glycoprotein I and II, while the lead compounds were not. However, all compounds were P-glycoprotein substrates.

**Table 1:** Chemical and physical properties of reference and lead compounds.

| Descriptors | OOU(reference) | Isopteropodin | Strophanthidin |
| --- | --- | --- | --- |
| Chemical formula | C19H19Cl2N3O | C21H24N2O4 | C23H32O6 |
| PubChem ID | 18353708 | 98363 | 6185 |
| Molecular Weight | 376.3 | 368.4 | 404.5 |
| (g/mol) |  |  |  |
| XLogP3 | 5 | 1.6 | 0.6 |
| HBD count | 3 | 1 | 3 |
| HBA count | 4 | 5 | 6 |
| Rotatable bond count | 7 | 2 | 2 |
| TPSA (Å<sup>2</sup>) | 56.92 | 67.9 | 104 |
| PAIN Alerts | None | None | None |
| Molar Refractivity | 105.6 | 106.47 | 106.16 |
| G-Protein CR Ligand | 0.31 | 0.37 | 0.08 |
| Ion channel modulator | 0.05 | 0.25 | 0.07 |
| Protein Kinase | 0.27 | -0.34 | -0.46 |
| Inhibitor |  |  |  |
| Nuclear Receptor | -0.2 | 0.07 | 0.52 |

| ligand |  |  |  |
| --- | --- | --- | --- |
| Protease Inhibitor | 0.1 | -0.02 | 0.01 |
| Enzyme Inhibitor | 0.24 | -0.02 | 0.79 |

**Table 2:**
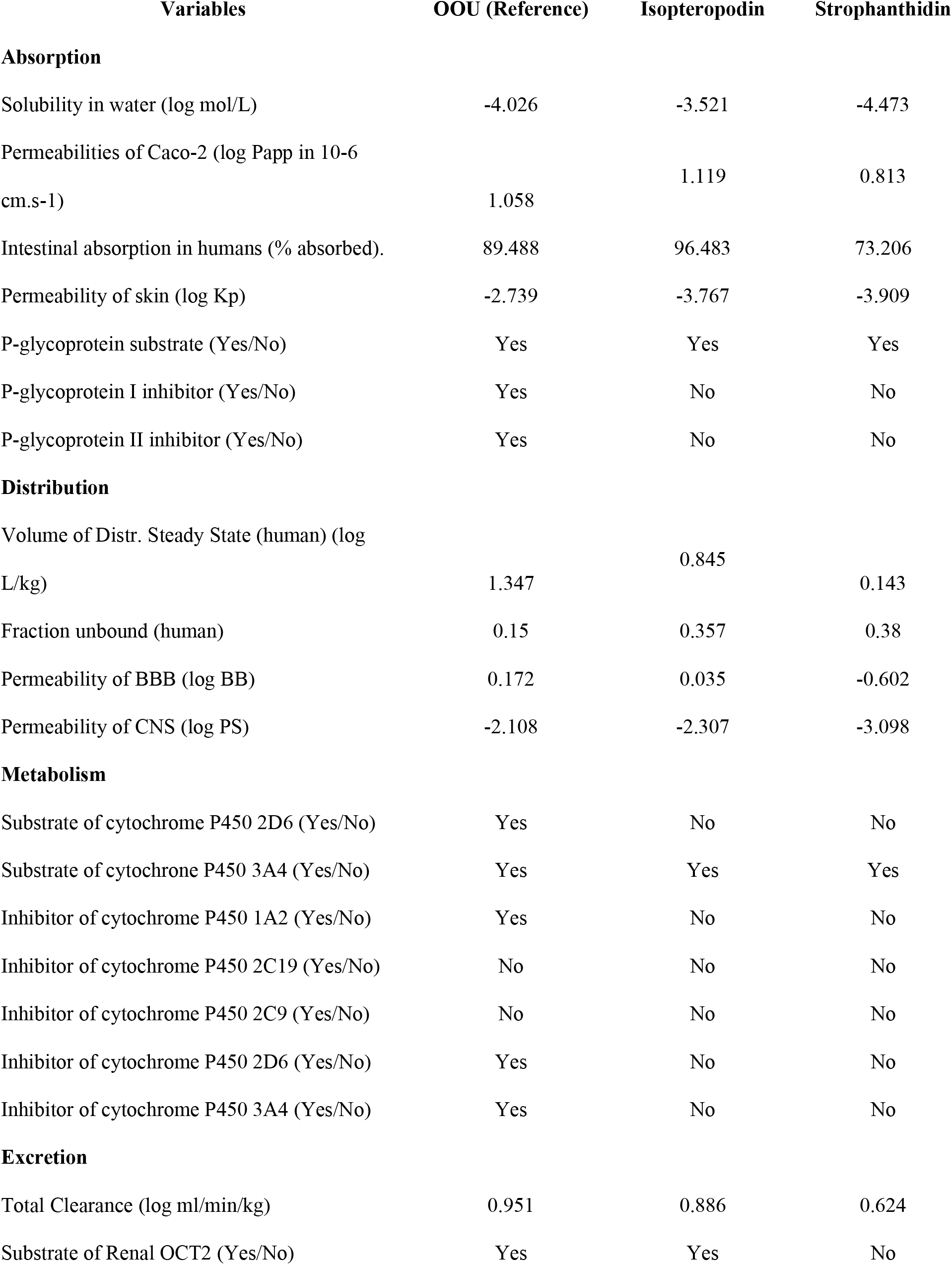
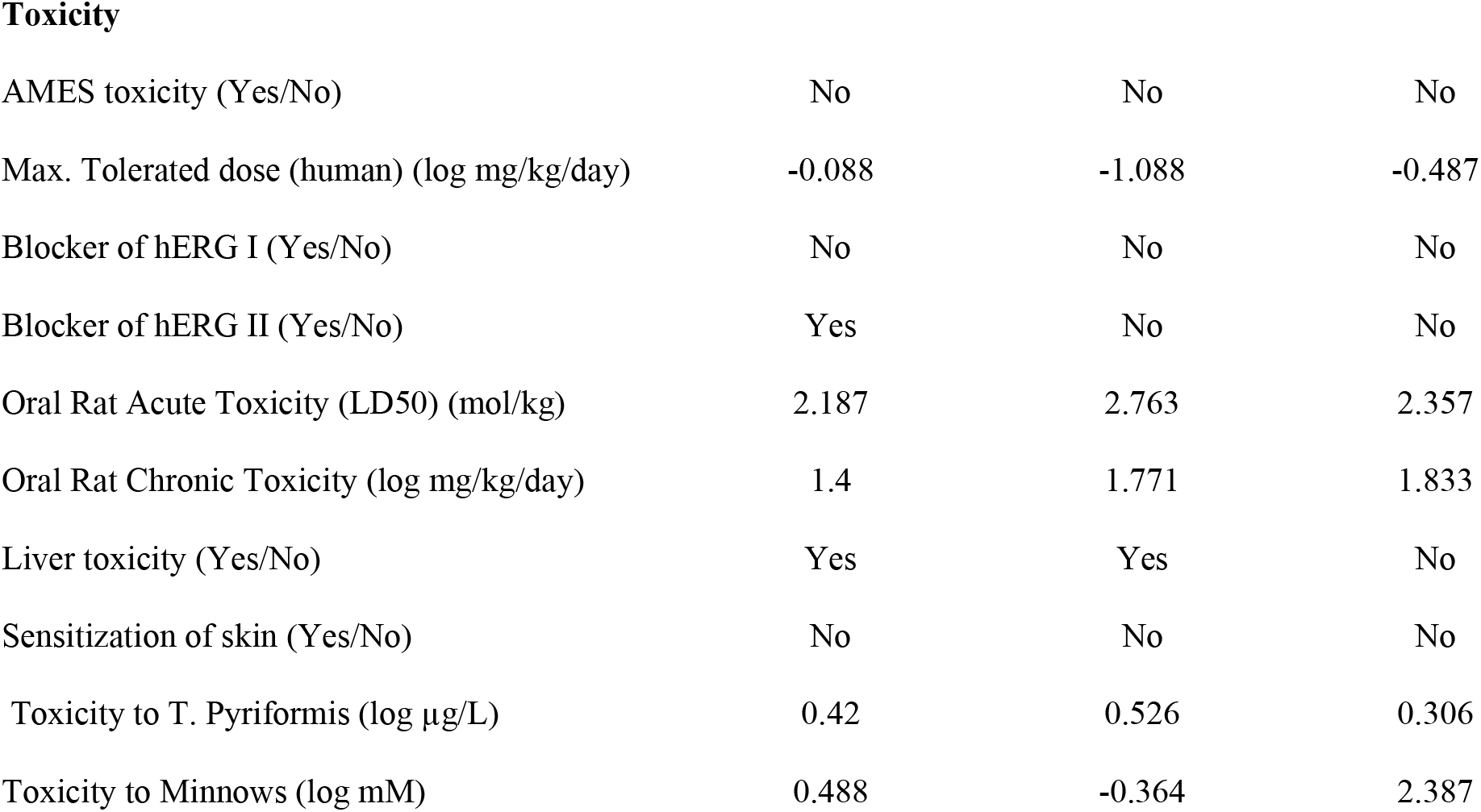
ADMET properties of reference and lead compounds.

| Variables | OOU (Reference) | Isopteropodin | Strophanthidin |
| --- | --- | --- | --- |
| **Absorption** |  |  |  |
| Solubility in water (log mol/L) | -4.026 | -3.521 | -4.473 |
| Permeabilities of Caco-2 (log Papp in 10<sup>-6</sup> cm.s<sup>-1</sup>) | 1.058 | 1.119 | 0.813 |
| Intestinal absorption in humans (% absorbed). | 89.488 | 96.483 | 73.206 |
| Permeability of skin (log Kp) | -2.739 | -3.767 | -3.909 |
| P-glycoprotein substrate (Yes/No) | Yes | Yes | Yes |
| P-glycoprotein I inhibitor (Yes/No) | Yes | No | No |
| P-glycoprotein II inhibitor (Yes/No) | Yes | No | No |
| **Distribution** |  |  |  |
| Volume of Distr. Steady State (human) (log L/kg) | 1.347 | 0.845 | 0.143 |
| Fraction unbound (human) | 0.15 | 0.357 | 0.38 |
| Permeability of BBB (log BB) | 0.172 | 0.035 | -0.602 |
| Permeability of CNS (log PS) | -2.108 | -2.307 | -3.098 |
| **Metabolism** |  |  |  |
| Substrate of cytochrome P450 2D6 (Yes/No) | Yes | No | No |
| Substrate of cytochrome P450 3A4 (Yes/No) | Yes | Yes | Yes |
| Inhibitor of cytochrome P450 1A2 (Yes/No) | Yes | No | No |
| Inhibitor of cytochrome P450 2C19 (Yes/No) | No | No | No |
| Inhibitor of cytochrome P450 2C9 (Yes/No) | No | No | No |
| Inhibitor of cytochrome P450 2D6 (Yes/No) | Yes | No | No |
| Inhibitor of cytochrome P450 3A4 (Yes/No) | Yes | No | No |
| **Excretion** |  |  |  |
| Total Clearance (log ml/min/kg) | 0.951 | 0.886 | 0.624 |
| Substrate of Renal OCT2 (Yes/No) | Yes | Yes | No |
| **Toxicity** |  |  |  |
| AMES toxicity (Yes/No) | No | No | No |
| Max. Tolerated dose (human) (log mg/kg/day) | -0.088 | -1.088 | -0.487 |
| Blocker of hERG I (Yes/No) | No | No | No |
| Blocker of hERG II (Yes/No) | Yes | No | No |
| Oral Rat Acute Toxicity (LD50) (mol/kg) | 2.187 | 2.763 | 2.357 |
| Oral Rat Chronic Toxicity (log mg/kg/day) | 1.4 | 1.771 | 1.833 |
| Liver toxicity (Yes/No) | Yes | Yes | No |
| Sensitization of skin (Yes/No) | No | No | No |
| Toxicity to T. Pyriformis (log µg/L) | 0.42 | 0.526 | 0.306 |
| Toxicity to Minnows (log mM) | 0.488 | -0.364 | 2.387 |

In terms of distribution, Strophanthidin had a CNS permeability (Log PS) value less than -3.0, while Isopteropodin and OOU had values larger than -3.0 but less than -2.0. The OOU and Isopteropodin had their volume of distribution steady state (Log VDss) values of more than 0.45, although Strophanthidin had a value of less than 0.15. All compounds had their BBB permeability (log BB) larger than -1.0 but less than 0.3. Similarly, all the compounds had their fraction unbound values greater than 0.1. With regards to metabolism, all compounds were non-inhibitors of cytochrome P450 2C19 and 2C9 enzymes and all substrates of cytochrome P450 3A4. Only OOU was an inhibitor of cytochrome P450 2D6, 1A2 and 3A4 enzymes and a substrate of cytochrome P450 2D6 (Table 2).

In terms of excretion, Strophanthidin recorded the lowest total clearance (log ml/min/kg), whereas OOU showed the highest. Strophanthidin was not a substrate of renal OCT2, only OOU and Isopteropodin were. All the compounds showed no AMES toxicity, no dermotoxocity and were non-inhibitors of hERG I proteins. However, only OOU was predicted to be a blocker of hERG II and only strophanthidin was not hepatotoxic. The values for maximum tolerated dose (log mg/kg/day), oral rat acute toxicity (LD50) (mol/kg) and oral rat chronic toxicity (log mg/kg/day) were highest in OOU, Isopteropodin, and strophanthidin respectively. For lead compounds and the reference, the *T. Pyriformis* toxicity (log g/L) values were all larger than - 0.5. Minnow toxicity (log mM) was less than 0.3 only for Isopteropodin (Table 2)

### Analysis of molecular docking scores

Isopteropodin had the lowest binding score with the target protein (Table 3).

**Table 3:** Docking scores of ligands against the target.

| Ligand | Binding Score (Kcal/mol) |
| --- | --- |
| OOU (reference) | -8.6 |
| Isopteropodin | -8.9 |
| Strophanthidin | -8.6 |

### Binding Site analyses

The reference and two lead compounds bound at residues ILE 12, TYR 14, VAL 229, TRP 230, ALA 233, LEU 234, GLY 262, ILE 265, PHE 268, PHE 293, and LEU 294 and could all be found in Pocket 1 of the target (Figure 5 & 6, Table 4 and supplementary data). The BrMelMetRS – Strophanthidin complex formed the highest number of intermolecular hydrogen bonds with the target. In terms of bond angle, Isopteropodin and Strophanthidin each formed one bond less than 130°at ILE12 and LEU294 respectively. The OOU, Isopteropodin, and Strophanthidin formed one, one and two bonds respectively that were greater than 130°. With reference to the donor to acceptor distance, OOU made no hydrogen bond within the range of 2.5-3.2 Å, none within the range of 3.2-4.0 Å, and only one (TYR14A) above 4.0 Å with the target. Isopteropodin formed one hydrogen bond (at ILE12A) within the range of 2.5-3.2 Å, and one (GLY262) within the range of 3.2-4.0 Å. Strophanthidin formed one hydrogen bond (at LEU294A) within the range of 2.5-3.2 Å, and two (at ILE21 and GLY262) within the range of 3.2-4.0 Å (Table 4). From Table 5, the BrMelMetRS – OOU complex had the highest number (12) of hydrophobic interactions and it was the only one that had a halogen bond at ASP232.

**Fig. 5:**
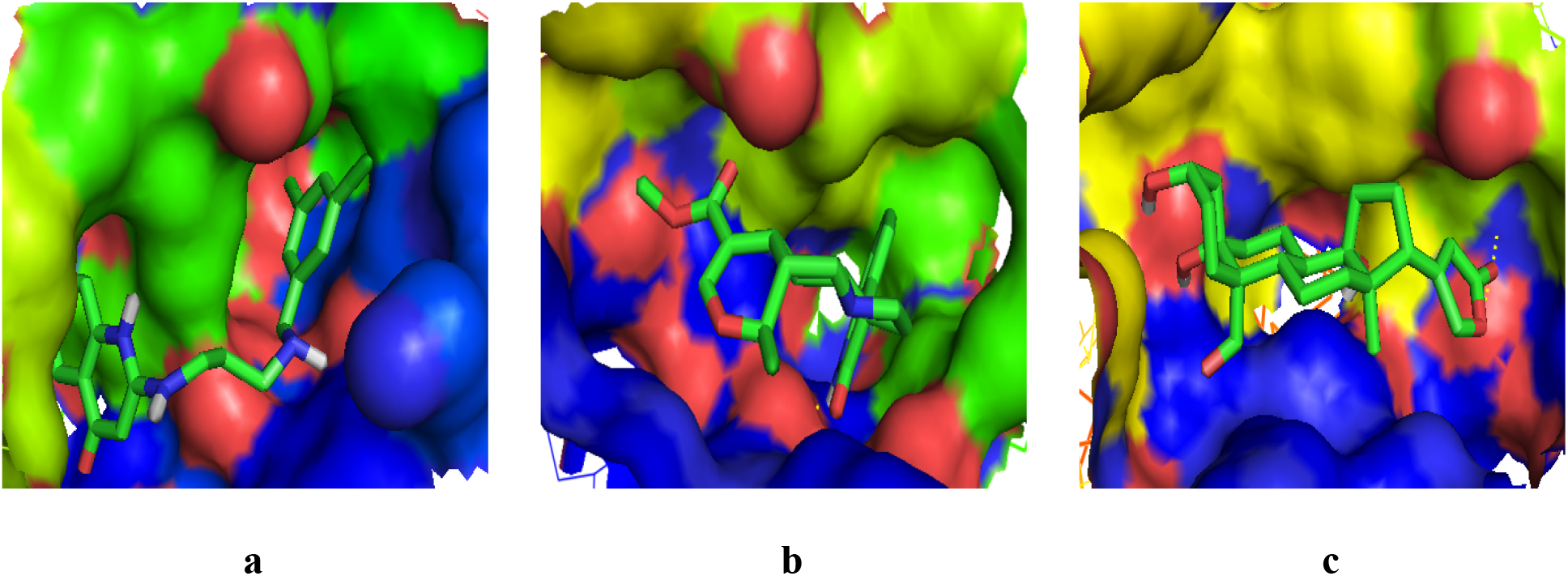
Binding site of the target showing interaction with reference lead compounds. (a) BrMelMetRS-OOU complex, (b) BrMelMetRS-Isopteropodin complex (c) BrMelMetRS-Strophanthidin complex

**Fig. 6:**
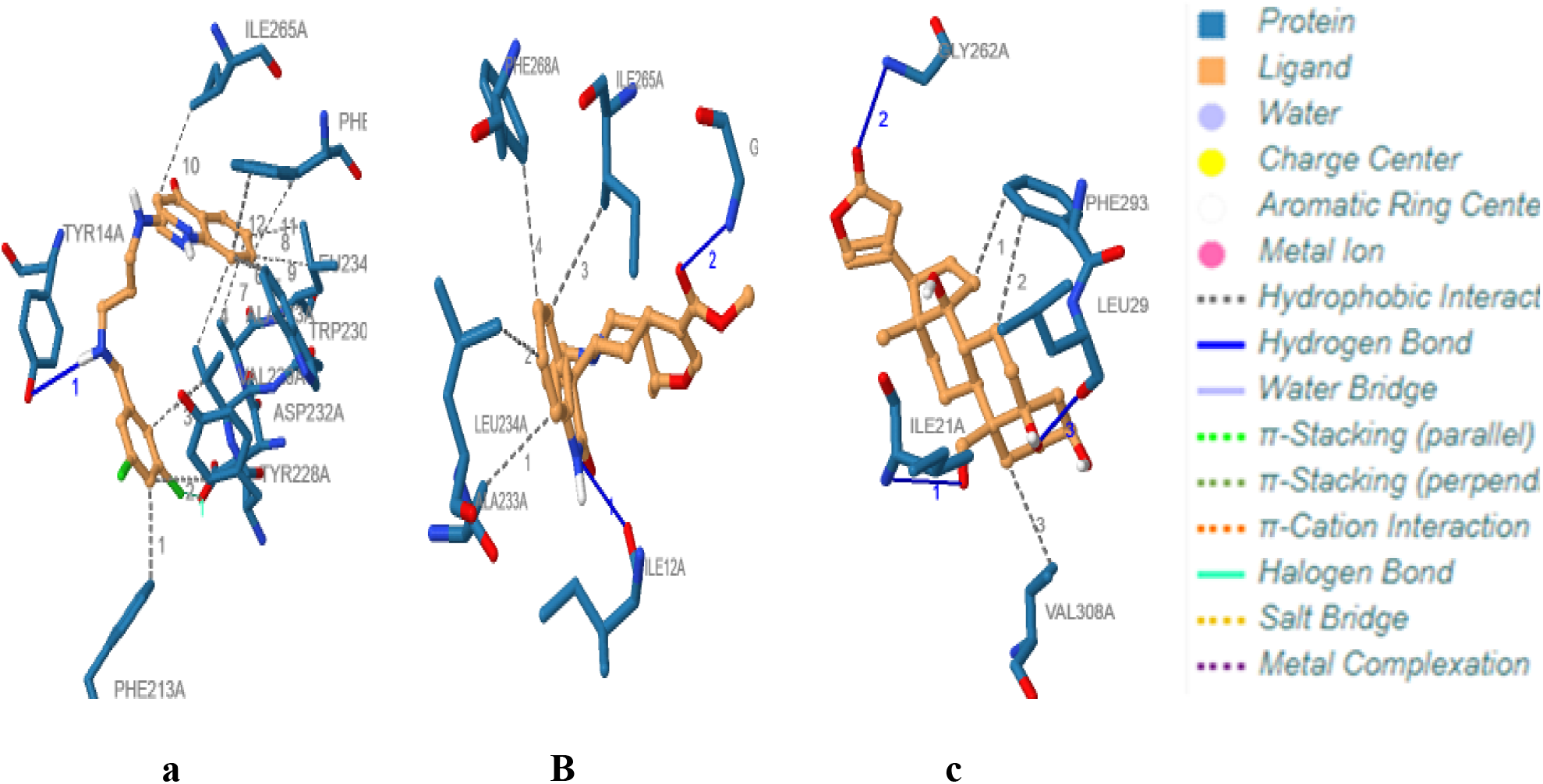
Interactions of target with reference and lead compounds (a) BrMelMetRS-OOU complex, (b) BrMelMetRS-Isopteropodin complex (c) BrMelMetRS-Strophanthidin complex

**Table 4:**
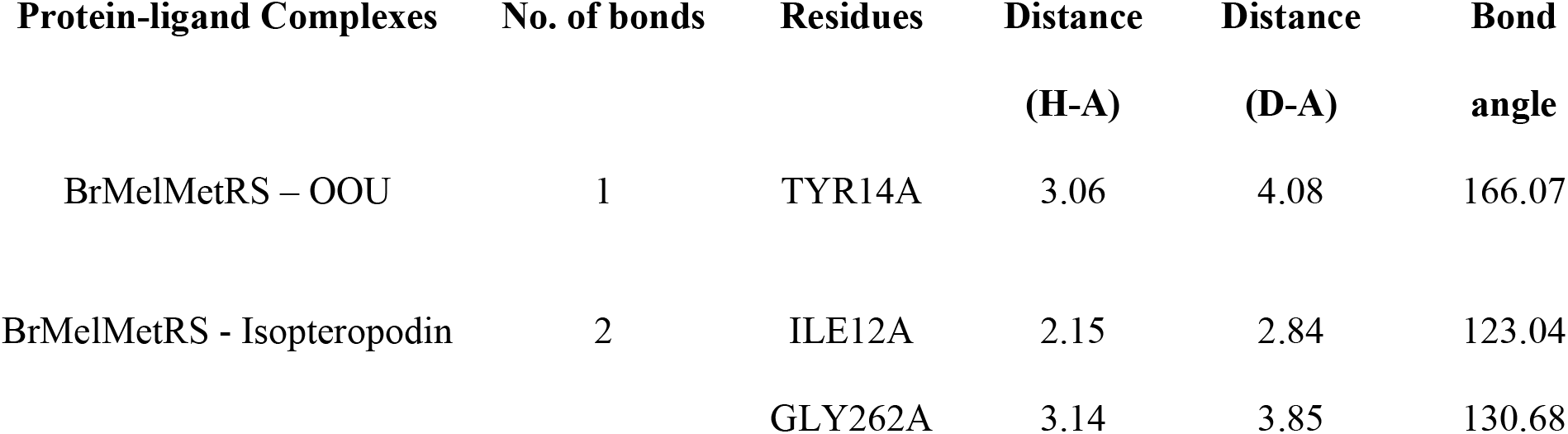
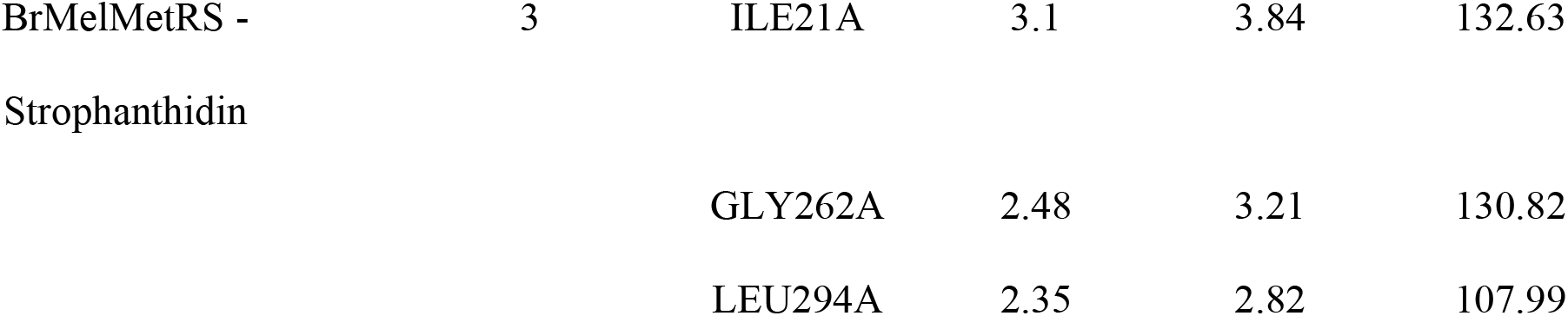
Analysis of Hydrogen bond interactions between target and ligands.

| Protein-ligand Complexes | No. of bonds | Residues | Distance | Distance | Bond angle |
| --- | --- | --- | --- | --- | --- |
|  |  |  | (H-A) | (D-A) |  |
| BrMelMetRS – OOU | 1 | TYR14A | 3.06 | 4.08 | 166.07 |
| BrMelMetRS - Isopteropodin | 2 | ILE12A | 2.15 | 2.84 | 123.04 |
|  |  | GLY262A | 3.14 | 3.85 | 130.68 |
| BrMelMetRS - | 3 | ILE21A | 3.1 | 3.84 | 132.63 |
| Strophanthidin |  | GLY262A | 2.48 | 3.21 | 130.82 |
|  |  | LEU294A | 2.35 | 2.82 | 107.99 |

**Table 5:**
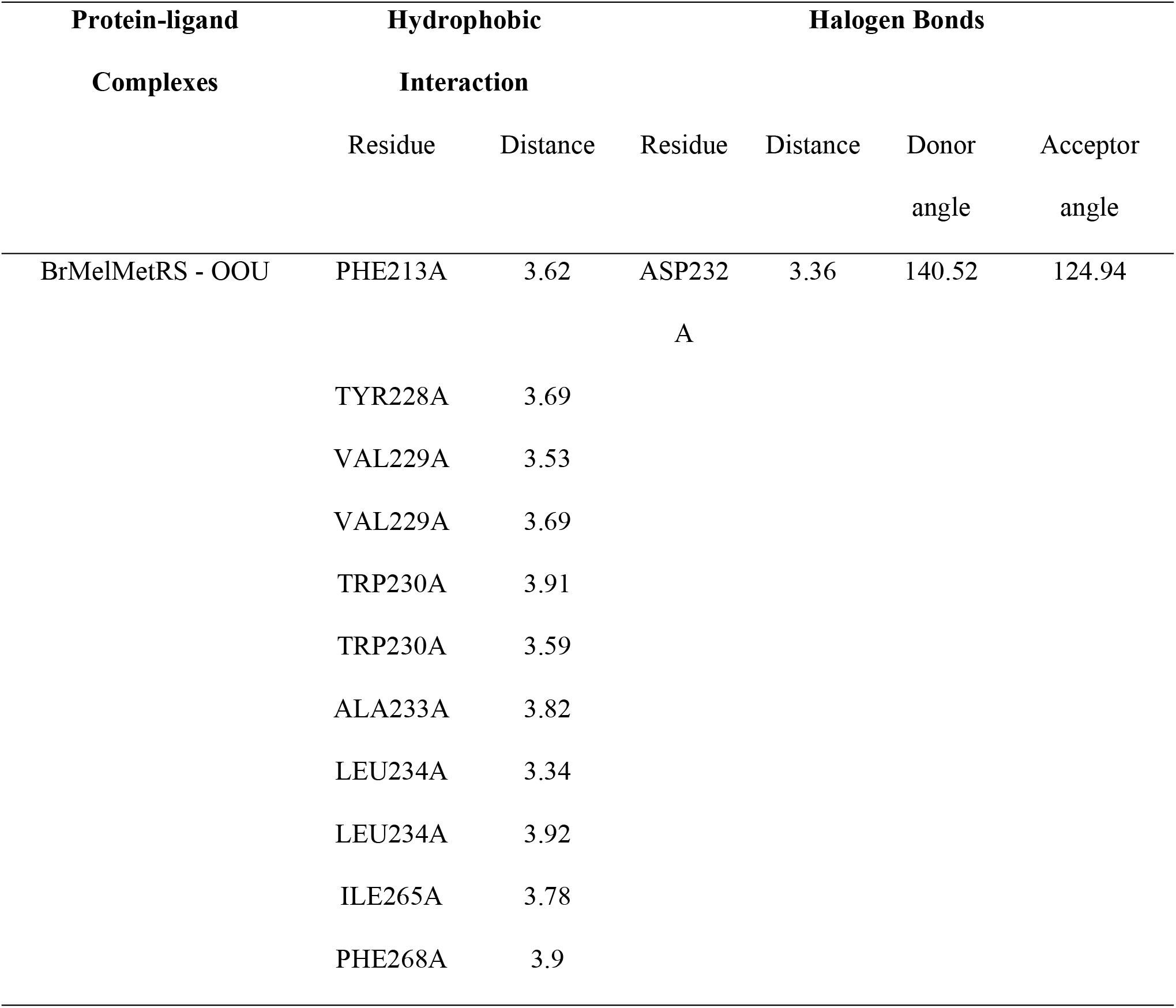
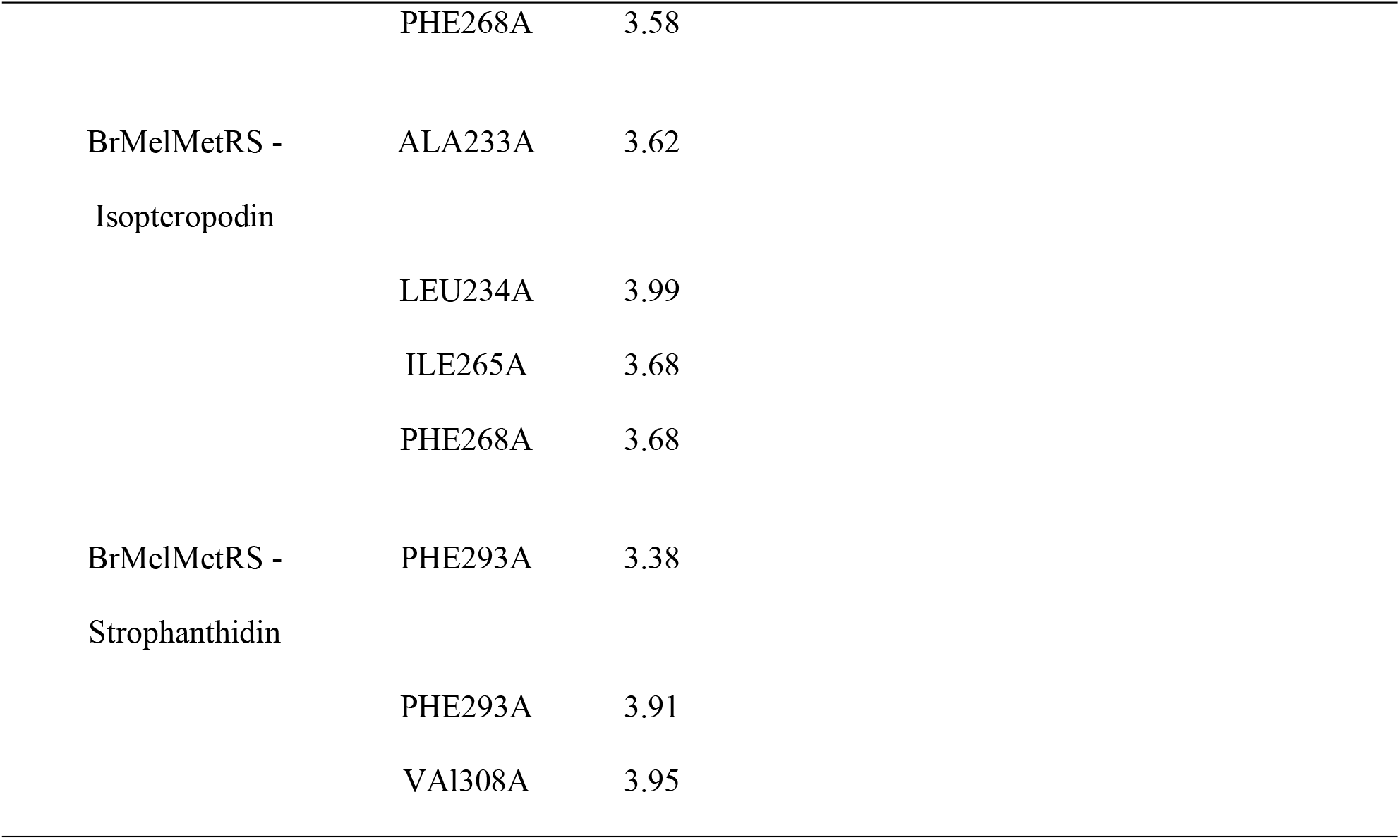
Hydrophobic interactions and Halogen bonds.

### Molecular dynamics simulation

Figure 7 reveals the structures of the apo and holo forms of BrMelMetRS after a 2-nanosecond molecular dynamics simulation. The BrMelMetRS-Isopteropodin and BrMelMetRS-OOU complexes had the same values of average and total RMSD which were higher than that of the BrMelMetRS-Strophanthidin complex. However, the BrMelMetRS-OOU trajectory peaked at time frame 15 (0.235) as compared with that of the BrMelMetRS-Isopteropodin complex which peaked at time frame 11 with a slightly lower RMSD value (0.231) (Figure 8 and Table 6). In terms of RMSD distribution, all the 20 peaks of the apo and holo forms of the target were found within 0.0 – 0.49 Å (Figure 9 and Table 6).

**Figure 7:**
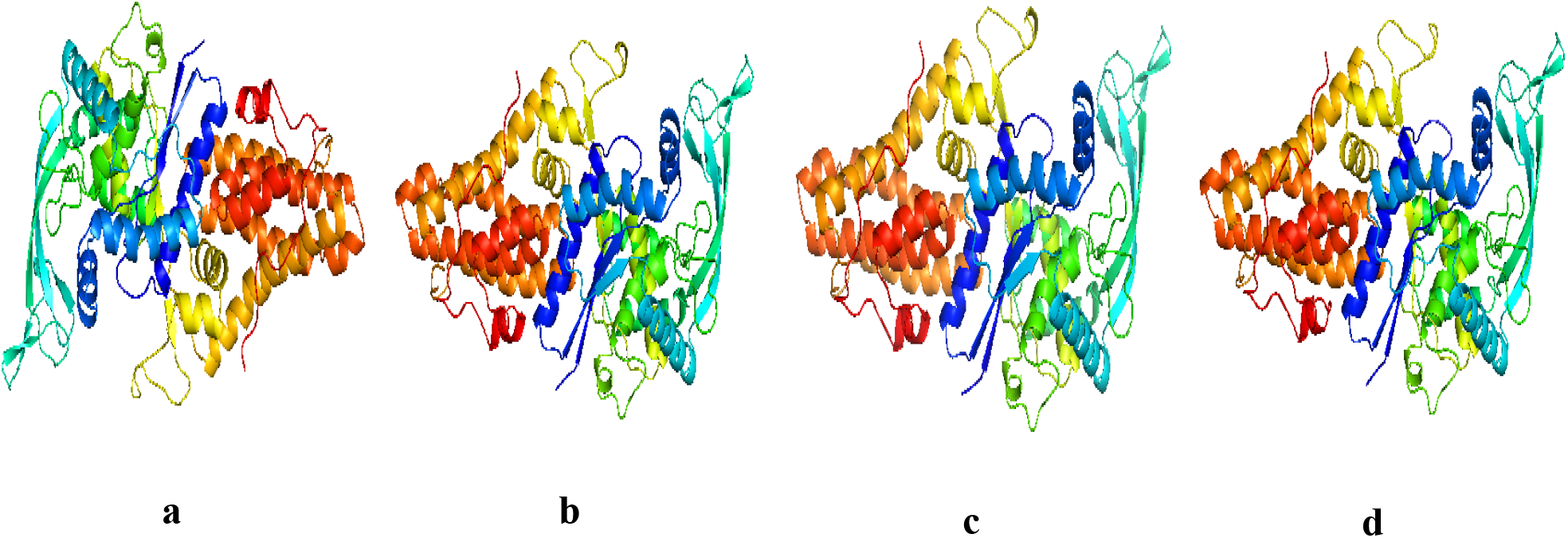
Cartoon model of the apo and holo forms of the target (after MDS). (a) BrMelMetRS (b) BrMelMetRS-OOU complex (c) BrMelMetRS-Isopteropodin complex (d) BrMelMetRS-Strophanthidin complex

**Figure 8:**
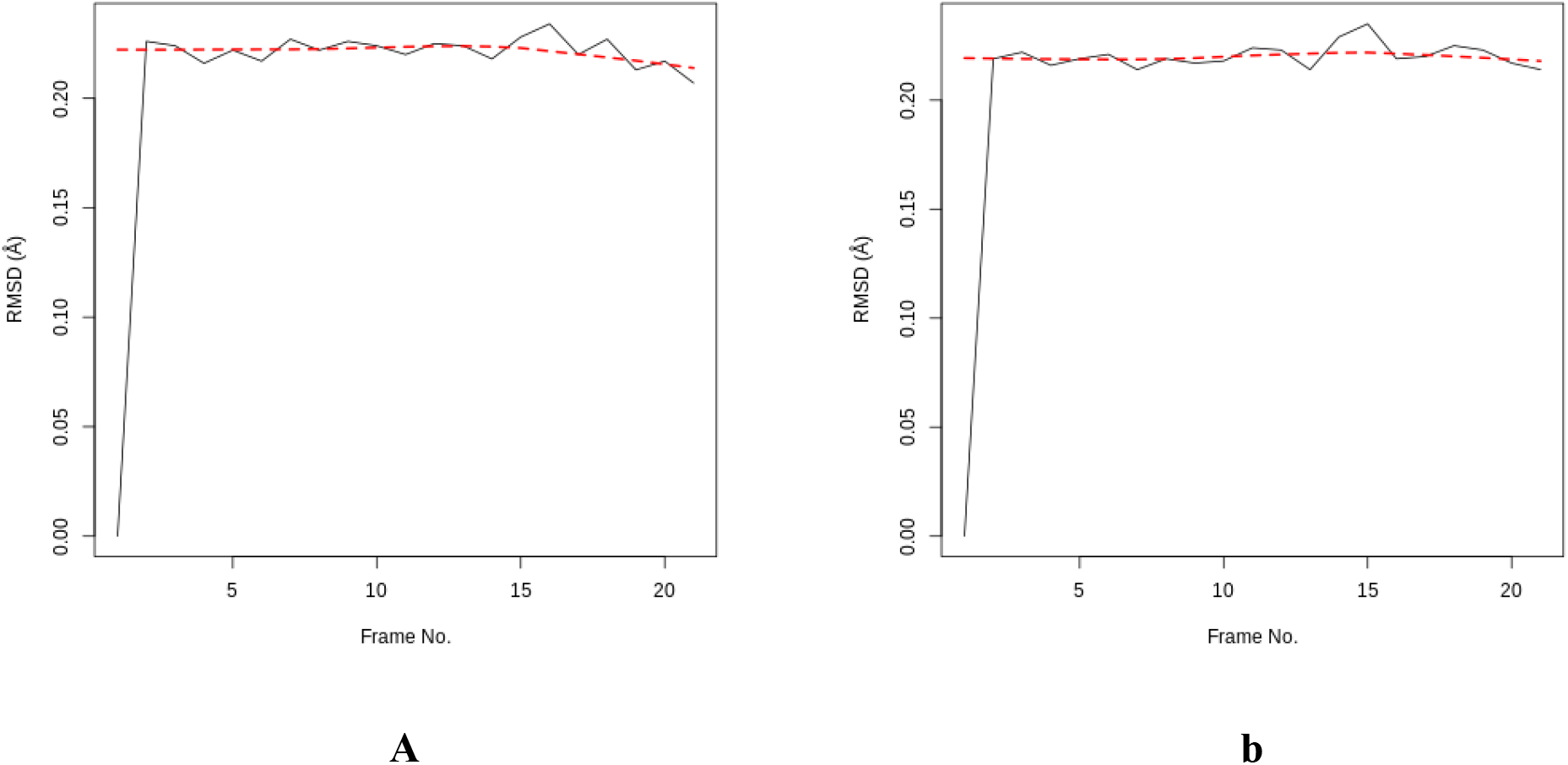
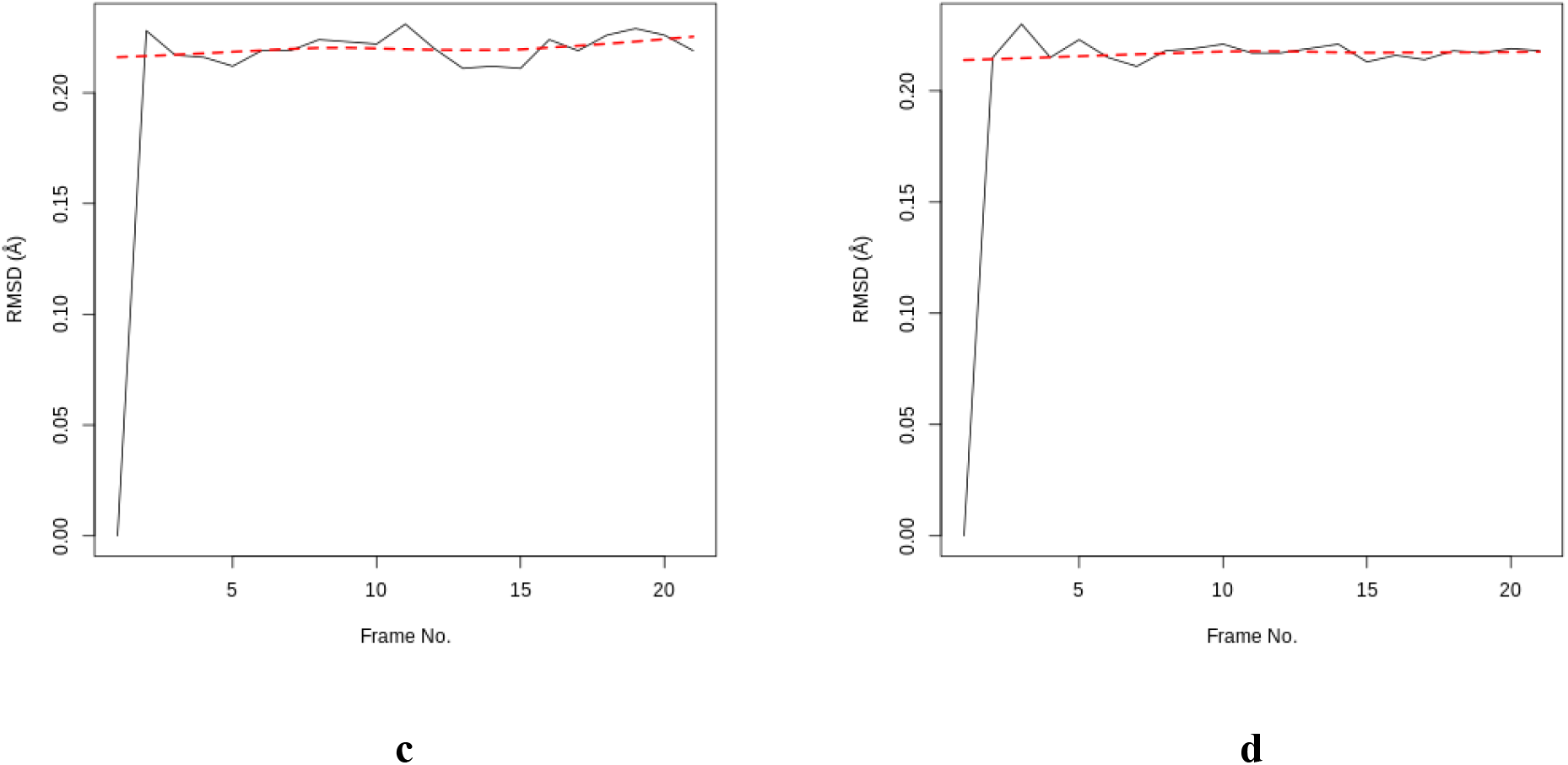
Root mean square deviations of the apo and holo forms of the target. (a) BrMelMetRS (b) BrMelMetRS-OOU complex (c) BrMelMetRS-Isopteropodin complex (d) BrMelMetRS-Strophanthidin complex

**Table 6:** MDS of the apo and holo forms of the target (a summary)

| MDS Parameters | BrMelMetRS<br>-Apo | BrMelMetRS<br>- OOU | BrMelMetRS -<br>Isopteropodin | BrMelMetRS -<br>Strophanthidin |
| --- | --- | --- | --- | --- |
| **RMSD** |  |  |  |  |
| Total RMSD | 4.437 | 4.408 | 4.408 | 4.356 |
| Average RMSD | 0.2112 | 0.2099 | 0.2099 | 0.2074 |
| Highest RMSD | 0.234 | 0.235 | 0.231 | 0.23 |
| Lowest RMSD | 0 | 0 | 0 | 0 |
| Time Frame of Highest RMSD | 16 | 15 | 11 | 3 |
| Time Frame of Lowest RMSD | 1 | 1 | 1 | 1 |
| RMSD Peak Distribution |  |  |  |  |
| 0.00 - 0.49A | 20 | 20 | 20 | 20 |
| 0.50 – 0.99A | 0 | 0 | 0 | 0 |
| 1.00 – 1.49A | 0 | 0 | 0 | 0 |
| 1.50 – 1.99A | 0 | 0 | 0 | 0 |
| 2.00 – 2.49A | 0 | 0 | 0 | 0 |
| 2.50 – 2.99A | 0 | 0 | 0 | 0 |
| 3.00 – 3.49A | 0 | 0 | 0 | 0 |
| **RMSF** |  |  |  |  |
| Total Global RMSF | 91.8492 | 91.5704 | 91.8067 | 91.0109 |
| Average Global RMSF | 0.1812 | 0.1806 | 0.1811 | 0.1795 |
| Total Regional RMSF(Pocket 1) | 5.4497 | 5.3549 | 5.3004 | 5.4541 |
| Average Regional RMSF(Pocket 1) | 0.1758 | 0.1727 | 0.171 | 0.1759 |
| Highest Fluctuation | 0.3149 | 0.2809 | 0.309 | 0.2884 |
| Least Fluctuation | 0.1038 | 0.1244 | 0.1183 | 0.122 |
| Range of RMSF | 0.2111 | 0.1565 | 0.1907 | 0.1664 |
| **PCA (motions)** |  |  |  |  |
| Total global (mean of PC1, PC2 & PC3) | 20.3527 | 20.5002 | 20.5907 | 20.4168 |
| Average global (mean of PC1, PC2 & PC3) | 0.0401 | 0.0404 | 0.0406 | 0.0403 |
| Total Regional (mean of PC1, PC2<br>& PC3) | 1.3102 | 1.2451 | 1.1877 | 1.2211 |
| Average Regional (mean of PC1,<br>PC2 & PC3) | 0.0423 | 0.0402 | 0.0383 | 0.0394 |
| Best global Conformation | PC3 | PC3 | PC2 | PC2 |
| Best regional Conformation<br>(Pocket 1) | PC3 | PC3 | PC2 | PC2 |
| PC1 Eigenvalue | 6.74 | 6.76 | 6.59 | 6.7 |
| PC2 Eigenvalue | 6.43 | 6.36 | 6.36 | 6.43 |
| PC3 Eigenvalue | 6.31 | 6.12 | 6.11 | 6.15 |
| Total | 19.48 | 19.24 | 19.06 | 19.28 |
| **B-Factor** |  |  |  |  |
| Global average B factor | 12.2297 | 27.7477 | 10.4732 | 15.6061 |
| Regional average B factor | 5.611 | 11.234 | 5.6169 | 7.1471 |
| **Radius of Gyration** |  |  |  |  |
| Average Gyration | 5.6951 | 5.6966 | 5.6951 | 5.6941 |
| Minimum Gyration | 5.693 | 5.6943 | 5.6908 | 5.6915 |
| Maximum Gyration | 5.6965 | 5.6984 | 5.6998 | 5.6966 |
| Range Gyration | 0.0035 | 0.0041 | 0.009 | 0.0051 |
| % Gyration | 0.061 | 0.072 | 0.158 | 0.09 |
| Time Frame of Maximum Gyration | 19 | 15 | 7 | 14 |
| Time Frame of Minimum Gyration | 13 | 12 | 3 | 20 |
| **DCCM** |  |  |  |  |
| Anti-correlating residues | 16 | 19 | 17 | 17 |

**Figure 9:**
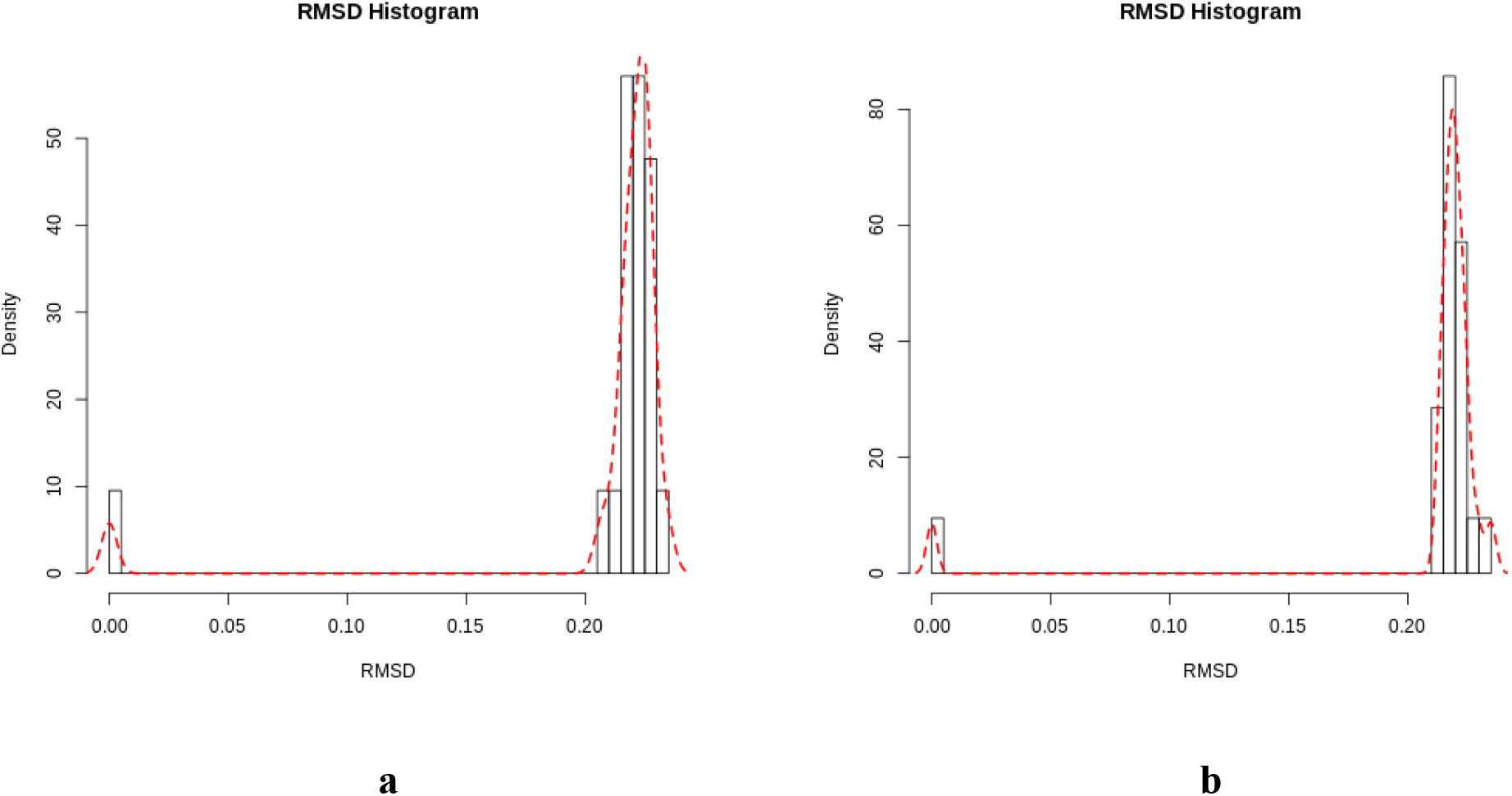
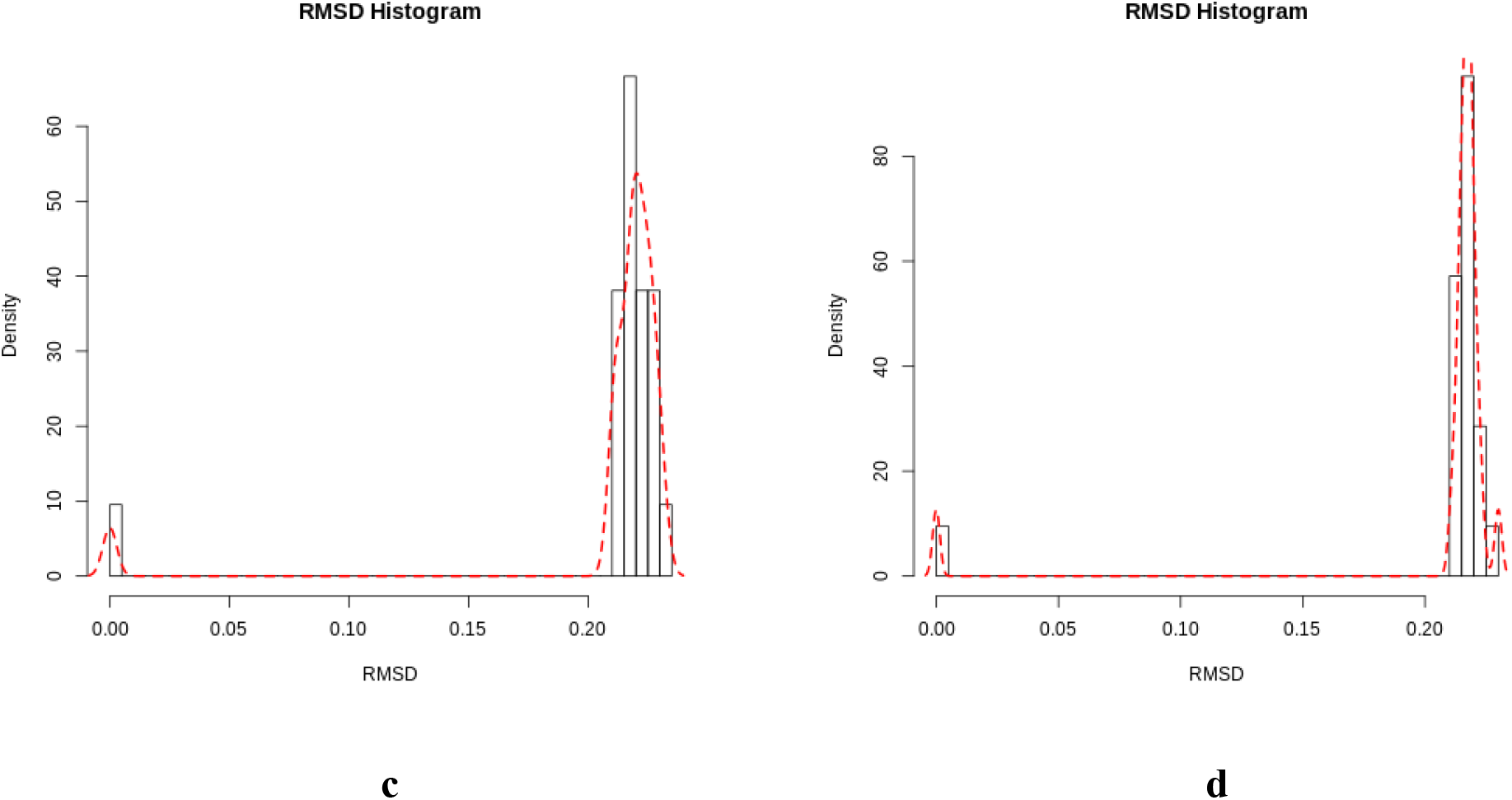
RMSD histogram of the apo and holo forms of the target. (a) BrMelMetRS (b) BrMelMetRS-OOU complex (c) BrMelMetRS-Isopteropodin complex (d) BrMelMetRS-Strophanthidin complex

The BrMelMetRS-Isopteropodin complex exhibited the greatest average and total RMSF values of all the holo structures. The least was the BrMelMetRS-Strophanthidin complex. However, at the regional level (Pocket 1), the BrMelMetRS-Strophanthidin complex had the highest values for average and total RMSF while that of the BrMelMetRS-Isopteropodin complex was the lowest (Figure 10 and Table 6).

**Figure 10:**
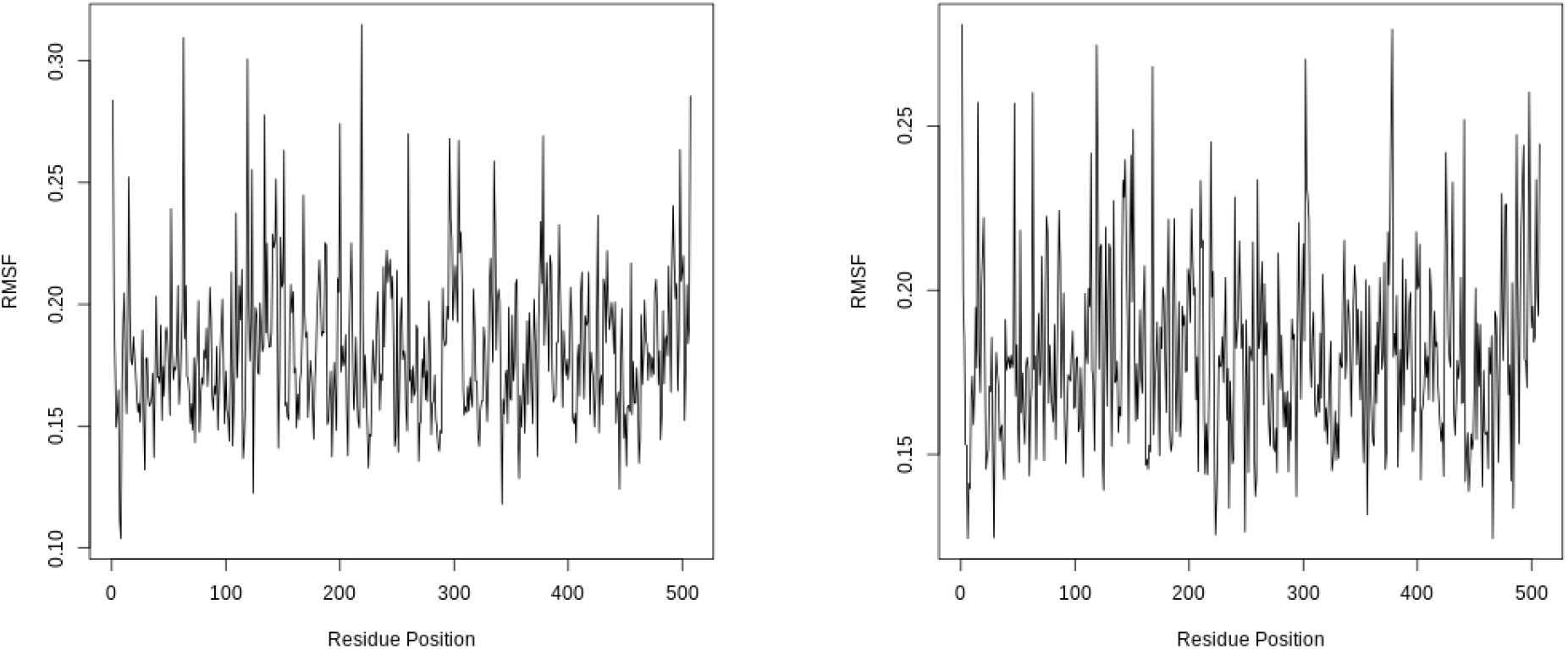
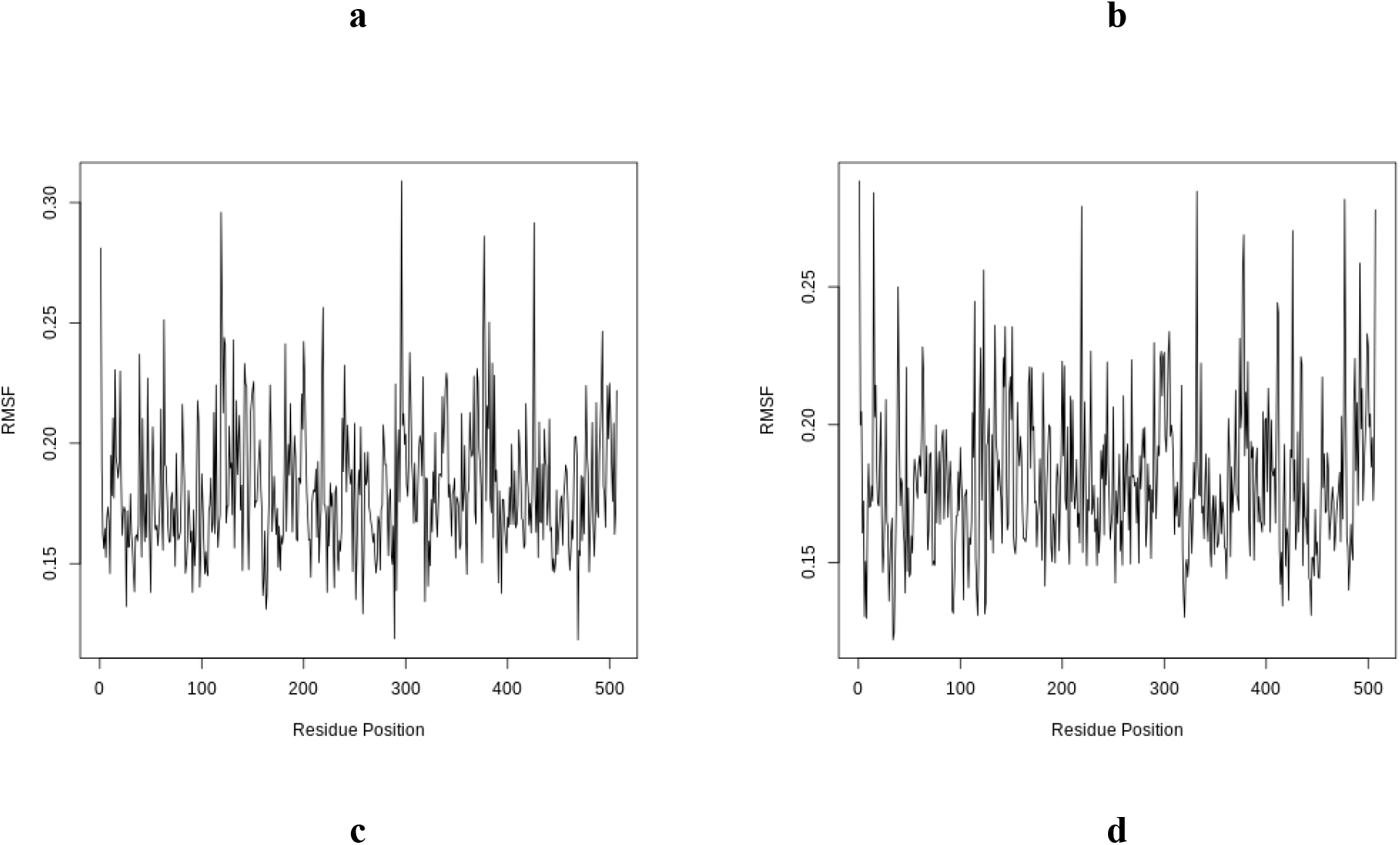
RMSF of the apo and holo forms of the target. (a) BrMelMetRS (b) BrMelMetRS-OOU complex (c) BrMelMetRS-Isopteropodin complex (d) BrMelMetRS-Strophanthidin complex

The cumulative of the first three highest principal components (PC1, PC2, and PC3) for all the holo forms of the target represented less than 50% of the total variance (Table 6 and Figure 11). The BrMelMetRS-Isopteropodin complex had the highest total and average global motions of all the holo forms. At the regional level (Pocket 1), the BrMelMetRS-OOU complex had the highest average and motions followed by the BrMelMetRS-Strophanthidin complex. Overall, the best conformations in terms of the greatest global motions were PC3, PC2, and PC2 for BrMelMetRS-OOU, BrMelMetRS-Isopteropodin complex, and BrMelMetRS-Strophanthidin complexes respectively and the same for regional motions. The PCA cosine content of the dominant motions related to PC1 for all the holo forms of the target did not get to 1.0 (Table 6).

**Figure 11:**
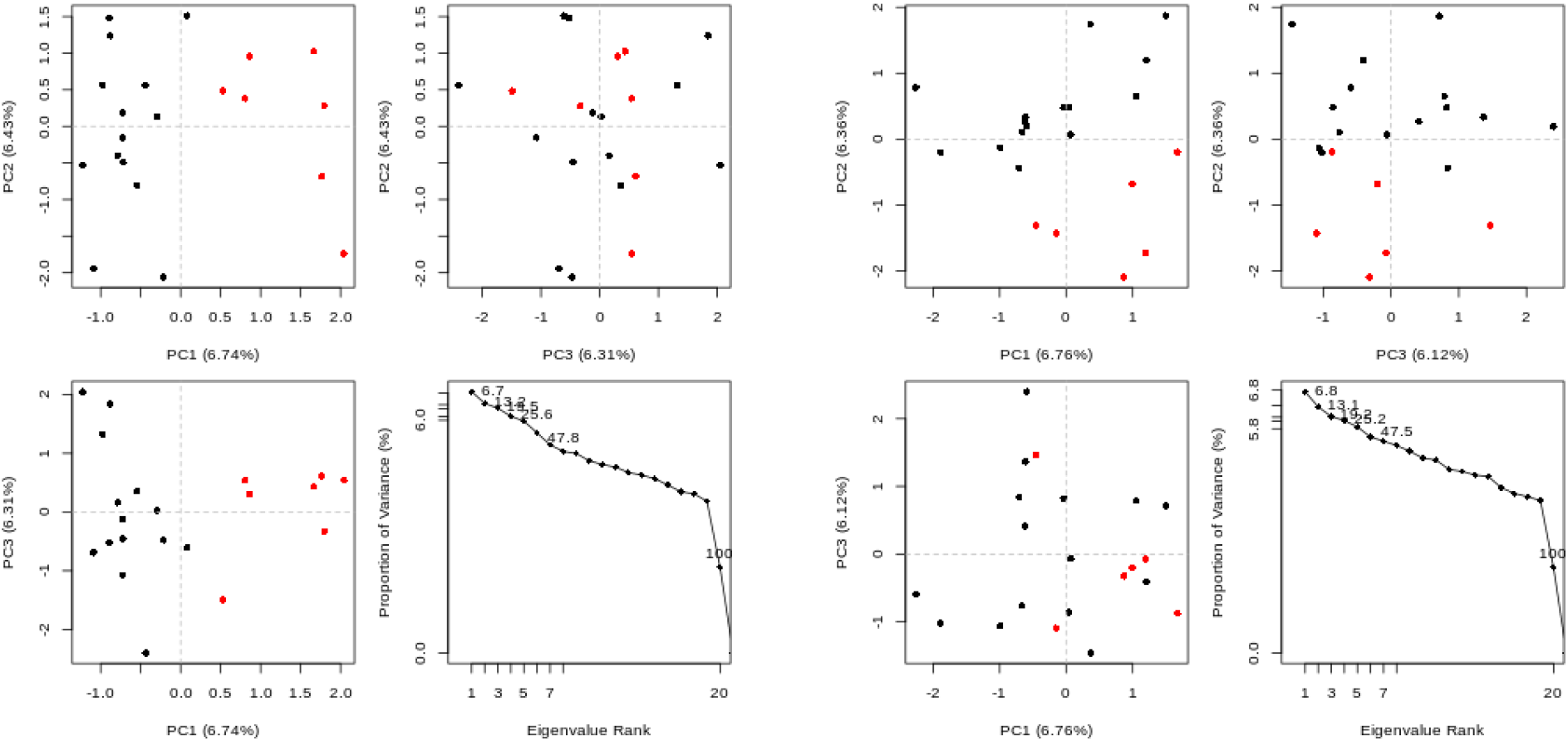
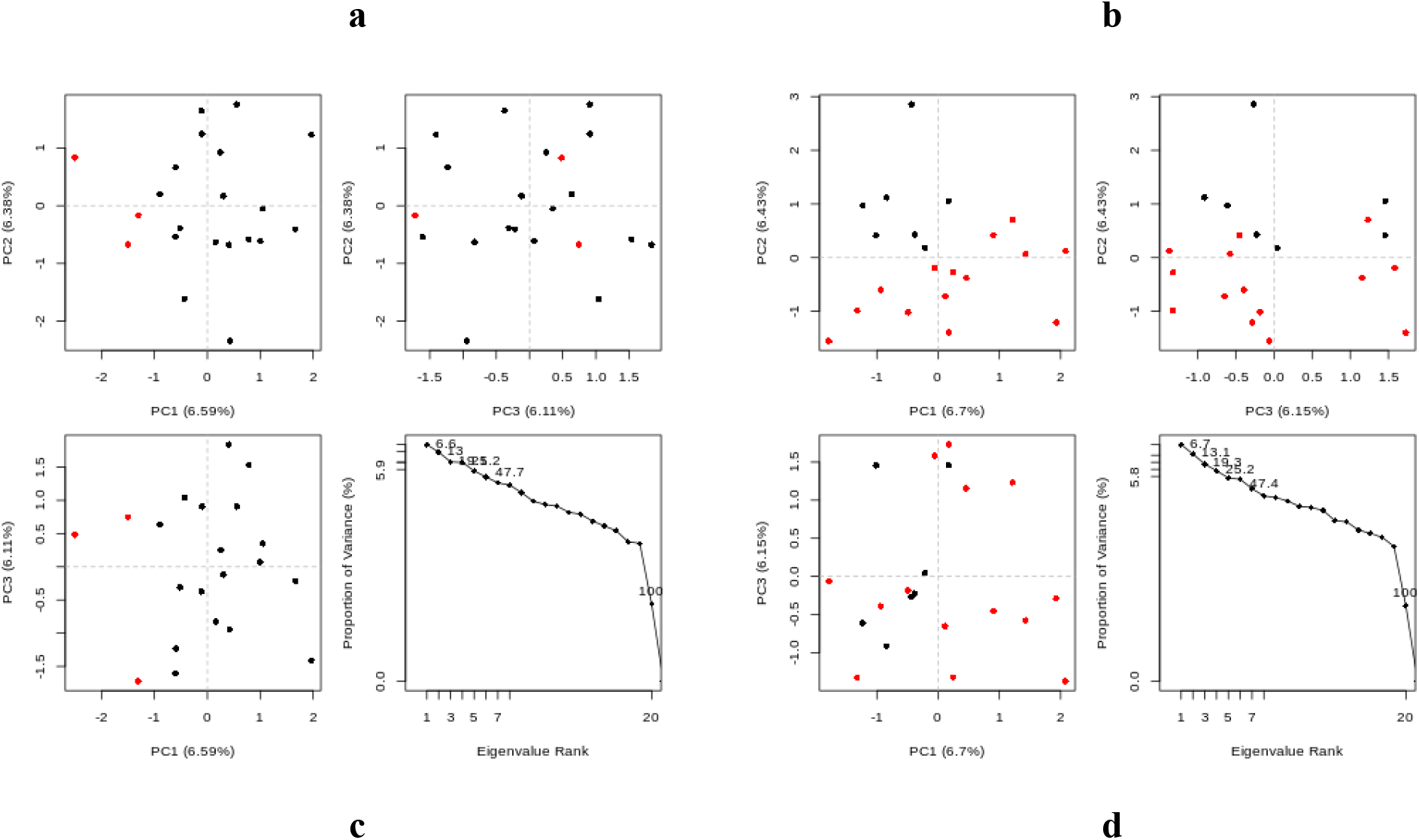
**PCA:** Cluster plots of the apo and holo forms of the target. The trajectory projection onto the first three eigenvectors for: (a) BrMelMetRS (b) BrMelMetRS-OOU complex (c) BrMelMetRS-Isopteropodin complex (d) BrMelMetRS-Strophanthidin complex

In terms of average radius of gyration along the trajectory, the BrMelMetRS-OOU complex had the highest value followed by the BrMelMetRS-Isopteropodin complex. However, the BrMelMetRS-Isopteropodin complex had the widest range of gyration (Figure 12 and Table 6). At the global and regional (Pocket 1) levels, B-factor values were highest in the BrMelMetRS-OOU complex (Figure 13 and Table 6). Additionally, the dynamic cross-correlation analysis revealed that of the 31 residues of the Pocket 1, the BrMelMetRS-OOU complex had the highest number of anti-correlating residues (Figure 14 and Table 6).

**Figure 12:**
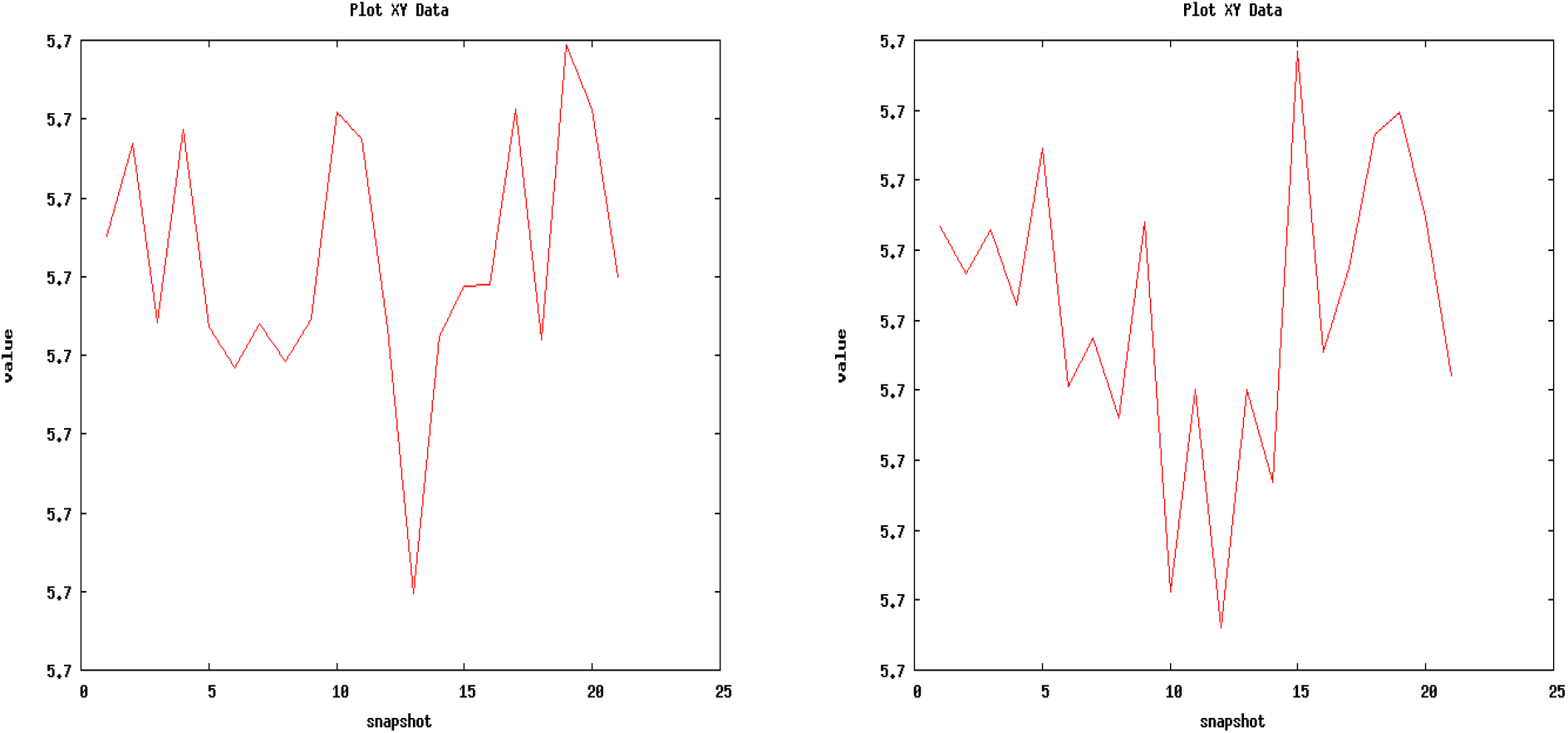
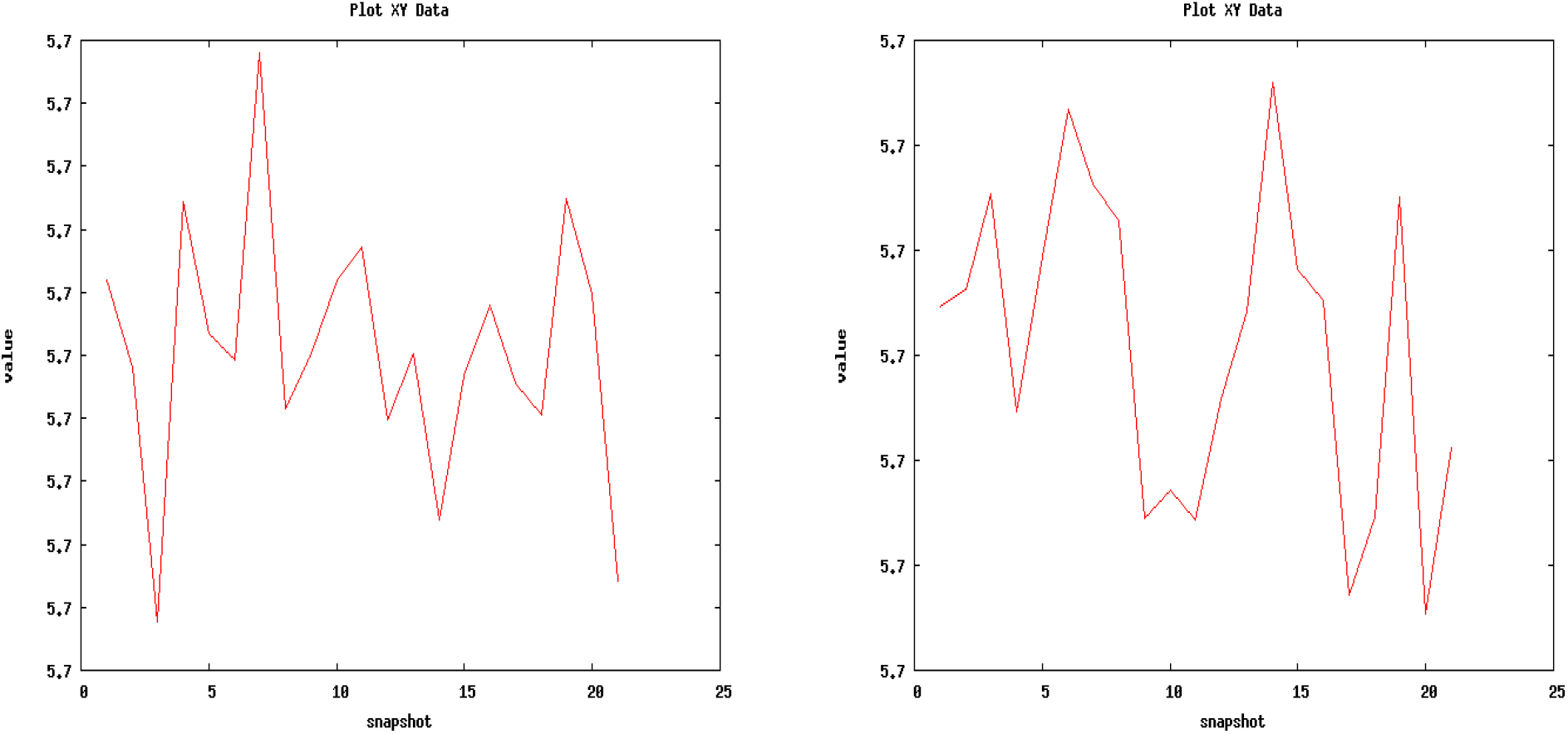
Radius of Gyration for the apo and holo forms of the target. (a) BrMelMetRS (b) BrMelMetRS-OOU complex (c) BrMelMetRS-Isopteropodin complex (d) BrMelMetRS-Strophanthidin complex

**Figure 13:**
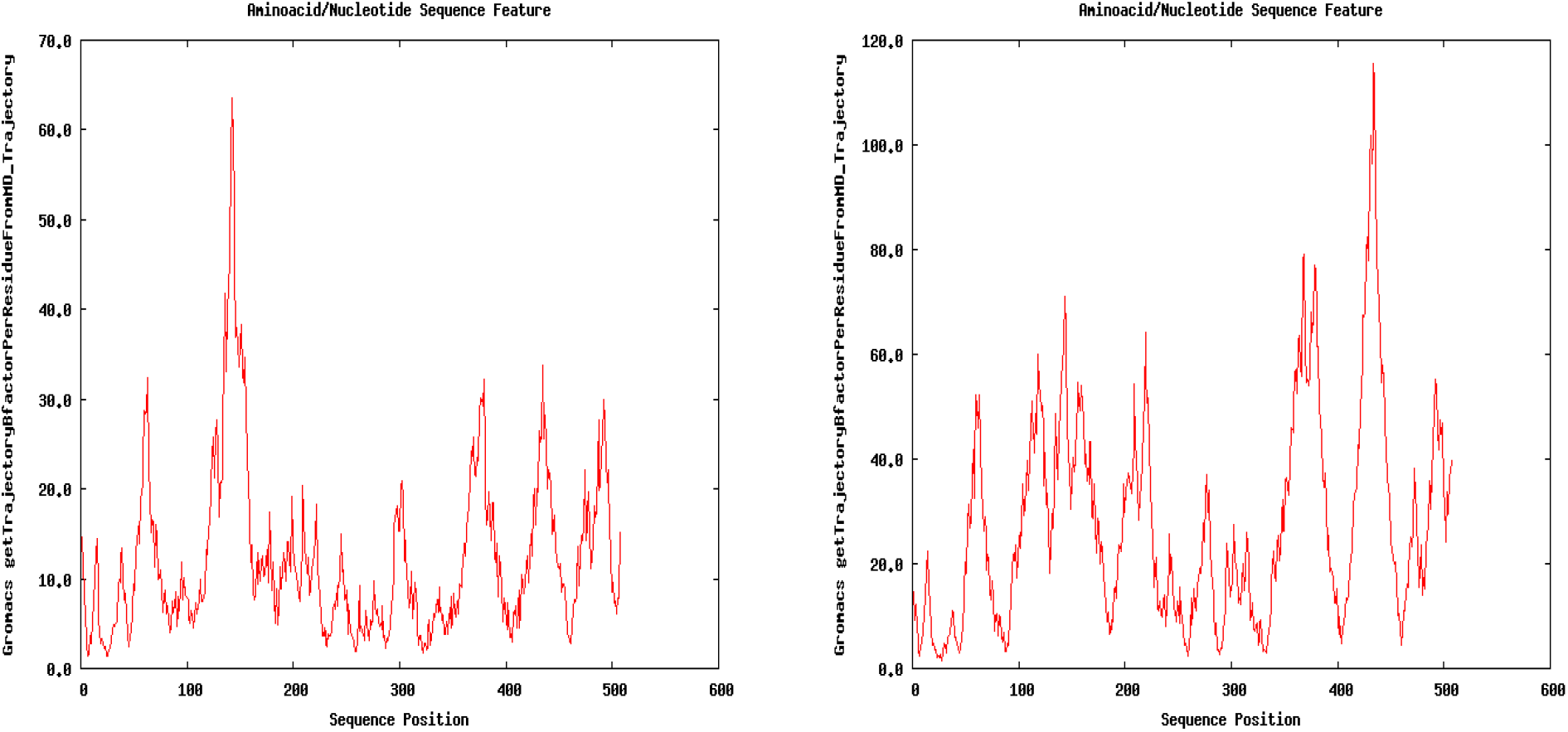
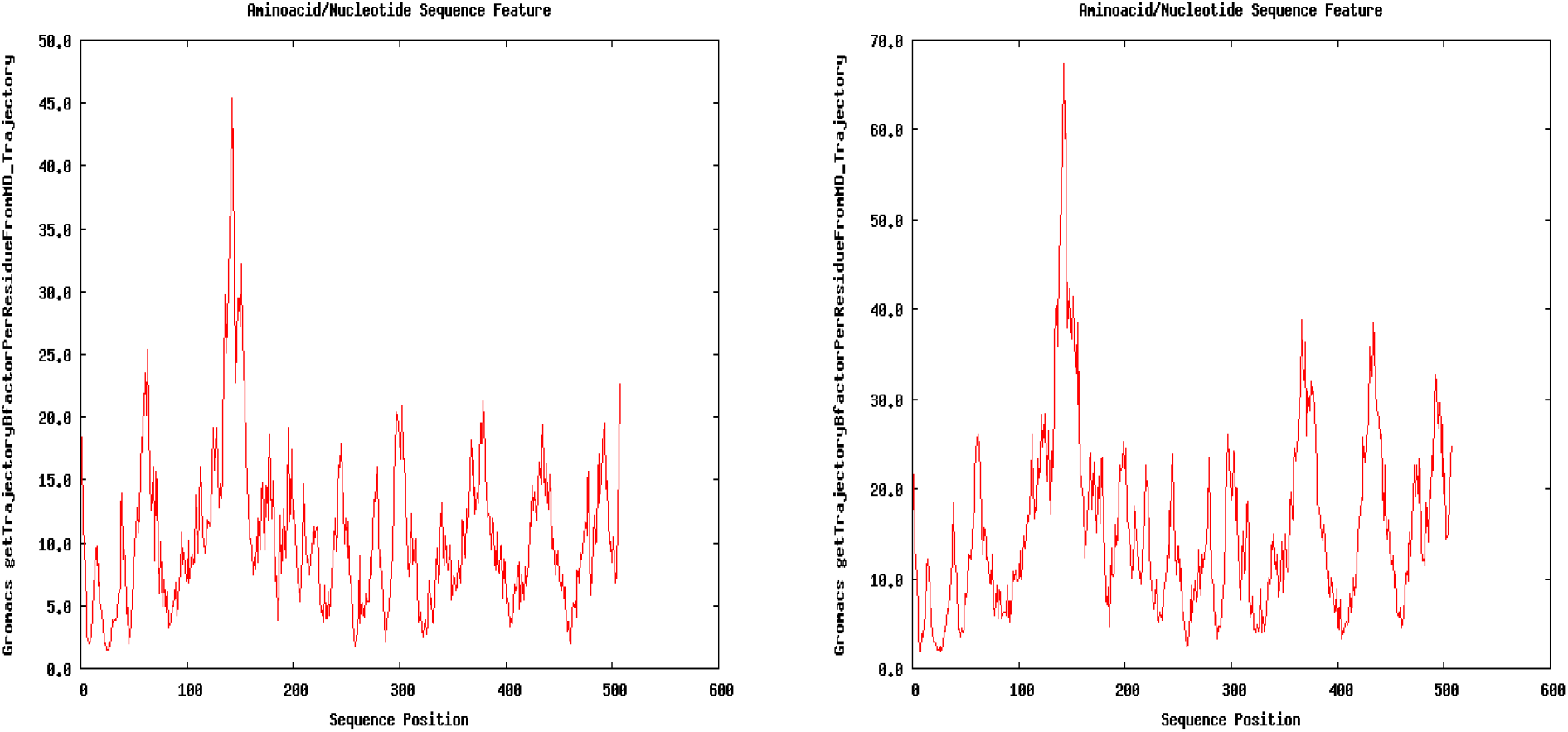
B-factor of the apo and holo forms of the target. (a) BrMelMetRS (b) BrMelMetRS-OOU complex (c) BrMelMetRS-Isopteropodin complex (d) BrMelMetRS-Strophanthidin complex

**Figure 14:**
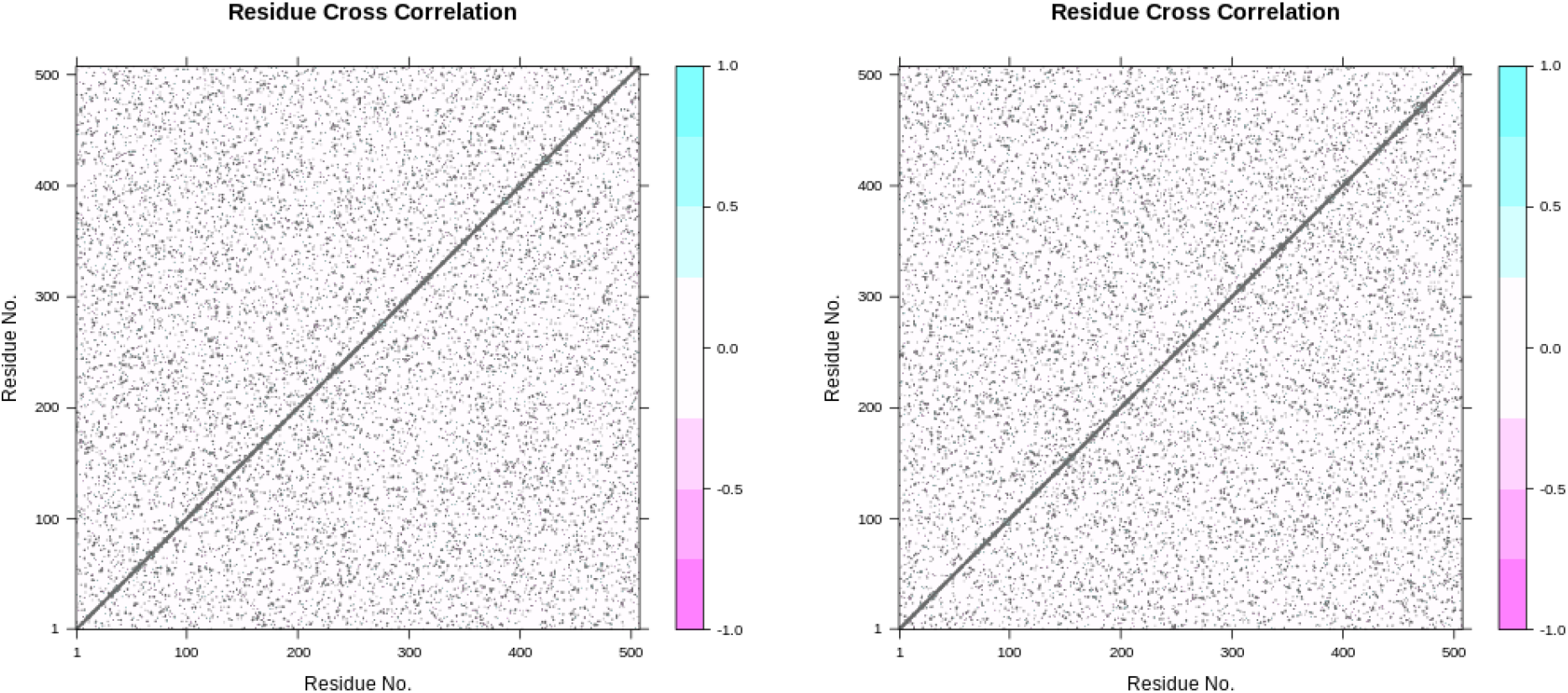
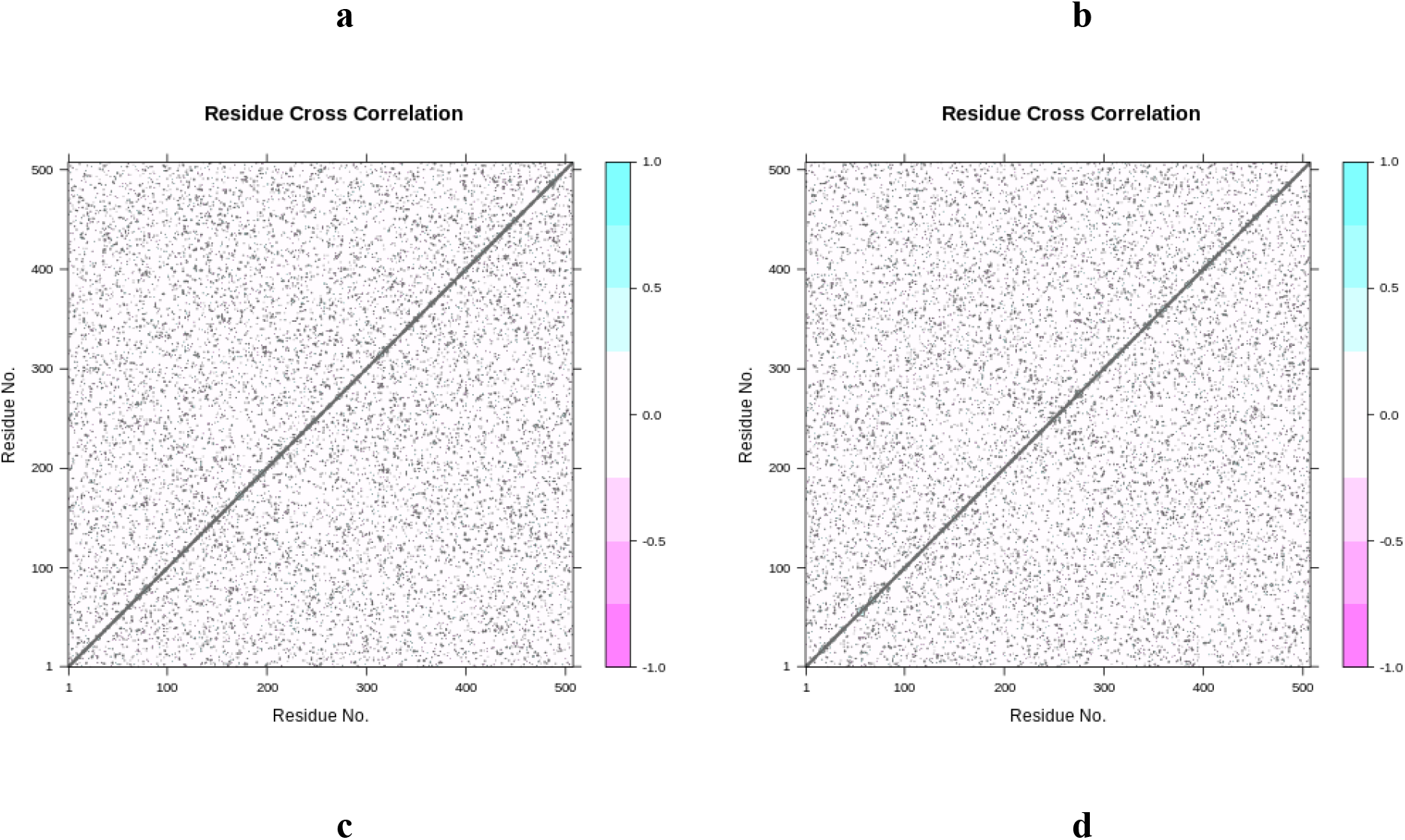
Dynamic cross correlation matrix of the apo and holo forms of the target. Dark cyan represents fully correlated motion, purple represents anti-correlated motion, while white and cyan represent moderately and uncorrelated motions respectively. Values of -1.0 are anti-correlated motion; 0 is non-correlated motion; and 1.0 is correlated motion. (a) BrMelMetRS (b) BrMelMetRS-OOU complex (c) BrMelMetRS-Isopteropodin complex (d) BrMelMetRS-Strophanthidin complex

### BLAST

The closest structures to the BrMelMetRS in the human proteome proteins were three unnamed protein products CBX51367.1, CAE90564.1 and CAE89160.1 (Table 7). The CBX51367.1 had a query cover of 91% while the other two showed short alignments each with 22% query cover.

**Table 7:** BLAST result for the homologues of the target protein in the human specie.

| Accession | Name | Accession | Max | Total | Query | E- | % |
| --- | --- | --- | --- | --- | --- | --- | --- |
|  |  | length | Score | Score | cover | value | identity |
| CBX51367 | unnamed protein | 900 | 148 | 148 | 91% | 4.00E- | 25.83% |
| .1 | product |  |  |  |  | 38 |  |
| CAE90564 | unnamed protein | 567 | 47 | 47 | 22% | 1.00E- | 34.43% |
| .1 | product |  |  |  |  | 05 |  |
| CAE89160 | unnamed protein | 764 | 45.1 | 45.1 | 22% | 5.00E- | 35.25% |
| .1 | product |  |  |  |  | 05 |  |

## Discussion

### The target

The structural quality of a protein determines its biological function [42]. The qualitative model energy analysis (QMEAN) is a composite scoring function assessing the major geometrical (global and local) aspects of *protein* structures. The quality assessment of the model which ranges from 0 to 1 (with one being perfect) is based on its distance from the template structure [43]. With a QMEAN value of 0.76, modelled BrMelMetRS (PDB: 5K0S.1.A) has a high structural quality. Similarly, the global model quality estimation (GMQE) score evaluates the structural quality of models using evolutionary information and it is expressed as a value between 0 and 1, with larger levels indicating greater reliability [44]. A value of 0.96 suggests a high reliability of the modelled target. Also, a ramanchandran favoured value greater than 98% and a Rama distribution Z-score less than 2 are suggestive of good stereochemistry of the modeled target [31].

### Drug-likeness and Bioactivity

A compound’s drug-likeness is determined by how similar it is to existing drugs in terms of structural and physicochemical features, and it serves as the foundation for future drug development [45]. In terms of drug-likeness, the reference and lead compounds do not violate the Ghose, Lipinski, and Veber rules seeing that the values of their HBA, HBD, log P, molecular weight, TPSA, molar refractivity, and number of rotatable bonds are within accepted range [46]. Therefore, they are all predicted to have good size, polarity and flexibility which positively correlate to good bioavailability. However, the bioavailability radar of OOU suggests that it is slightly unsaturated with fraction of carbons in the sp3 hybridization (Fsp^3^) value less than 0.25 [47,48]. Both complexity (as assessed by Fsp^3^) and the presence of chiral centers have been shown to be predictive of success as compounds go from discovery to clinical testing to drug development. It has also been shown that saturation is linked to solubility, which is crucial in drug development [47]. Therefore, OOU would be a poor drug candidate.

The development and appropriate use of new medications necessitate an examination of their biological activity profiles, which must take into account human metabolism. This is due to the fact that most drugs interact with several or multiple molecular targets in the body, resulting in complicated bioactivity profiles [49]. With respect to bioactivity, Strophanthidin is predicted to have the highest enzyme inhibition activity while Isopteropodin has the lowest (Table 1).

### ADMET

The ADMET properties of candidate compounds are the main reason of high attrition rates in drug discovery [50]. In the research and development of drugs, aqueous solubility is a critical physicochemical feature that influences pharmacokinetic properties and formulations [51]. From Table 2, Isopteropodin is the most soluble of the compounds. With water solubility values less than -4.0 log mol/L, OOU and Strophanthidin are poorly soluble [35]. Oral administration remains the primary method of administration, making *in vitro* permeability studies useful for predicting oral bioavailability throughout drug development. The Caco-2 cell monolayers, which produce tight connections between cells, are employed as a model of human intestinal absorption because they closely resemble the human intestinal epithelium in many ways [52]. Isopteropodin showed the highest Caco-2 permeability. The OOU also has high Caco-2 permeability while that of Strophanthidin is low [35]. Similar to Caco-2 permeability, the determination of human intestinal absorption (HIA) is a very important aspect in the creation of novel pharmacological compounds [53]. Though all the compounds have high percentage HIA, Isopteropodin has the highest value [35]. In order to build a transdermal medication delivery system for human usage, it is necessary to assess drug penetration through the skin [54]. While all the compounds had skin permeability (LogKp) values less than 2.5, Strophanthidin has the best dermal permeability value.

Expressed in the cells of several organs, ATP-binding cassette (ABC) superfamily of transporter proteins, P-glycoprotein (Pgp) has an influence on the ADMET properties of drugs. The Pgp is a unidirectional efflux pump that removes its substrate such as drugs, pollutants, and other xenobiotics from inside to outside of the cells [55]. All the compounds are P-glycoprotein substrates. This implies that their oral bioavailability would be reduced by P-glycoprotein. Unlike the lead compounds, OOU is predicted to be P-glycoprotein I and II inhibitors suggesting that it would facilitate the intracellular accumulation of substrates leading to toxicity [56].

The volume of distribution steady state (VDSS) is the theoretical volume that would be necessary to hold the whole amount of an administered medication at the same concentration that it is detected in the blood plasma. It is an important pharmacokinetic property that determines the dosing frequency and half-life of a drug [57]. The VDSS for OOU is extremely high requiring about 8.41l/kg; the VDSS of Isopteropodin is high requiring about 5.28 l/kg; and the VDSS of Strophanthidin is low requiring about 0.89 l/kg to maintain uniform distribution to give the same concentration in the plasma [35]. The degree to which a drug binds to plasma proteins has an impact on its efficacy. The drug can penetrate the cellular membranes more efficiently with less binding [35]. Though they all exceeded 0.1, the values of fraction unbound (human) for OOU suggests that it is the least available for bioactivity. Though Isopteropodin and Strophanthidin have similar values, Strophanthidin is more available [35]. The anatomically and functionally distinct blood–brain barrier (BBB) prevents the uptake of most drugs. However, certain drugs with unique chemical properties are able to cross the BBB through lipid-mediated free diffusion [58]. From Table 2, all compounds have their BBB permeability (log BB) larger than -1.0 but less than 0.3 suggesting that they are all moderately distributed in the brain. The OOU is predicted to have the best brain distribution while Strophanthidin has the poorest [35]. Similarly, Strophanthidin is also unable to permeate the CNS, while Isopteropodin and OOU can moderately permeate [35].

Cytochromes P450 (CYP) is responsible for the biotransformation of most medicines in clinical use and is a primary cause of variability in drug pharmacokinetics. The CYPs 3A4, 2C9, and 1A2 are the most prevalent in the liver, while 2D6 and 2C19 are less abundant [59]. All compounds are substrates of CYP450 3A4 and only OOU is a substrate of CYP450 2D6. This suggests that these metabolic enzymes facilitate the biotransformation of these compounds making them available for excretion. Remarkably, OOU is an inhibitor of CYP450, P450 2D6, 1A2, and 3A4 causing the accumulation of the substrate of these enzymes [35].

The total clearance of a drug from the bloodstream is the sum of the renal clearance, the hepatic clearance, and the clearance from all other tissues [60]. Depending on the functionality of the organs involved and several other factors, the total clearance ranges from 0 to 1.0. The results as indicated in Table 2 show that OOU followed by Isopteropodin have a very high rate of elimination from the plasma while that of Strophanthidin is slowest.

The renal organic cation transporter 2 (OCT2) protein is found in the basolateral membrane of proximal epithelial cells and it is involved in cationic drug uptake and secretion [61]. Only Strophanthidin, as shown in Table 2, will not be carried from the plasma into the cells of the proximal convoluted tubule by the renal OCT2 and will also have no deleterious interactions when co-administered with renal OCT2 inhibitors [35].

The potassium channel protein expressed by the human ether-a-go-go related gene (hERG) is important for cardiac repolarization and arrhythmias caused by long QT wave [62]. The study also found that only OOU is predicted to be an inhibitor of hERG II protein showing its potential cardiotoxic property [35]. However, all the compounds are neither genotoxic nor dermato-toxic. As established by early-stage human clinical trials, the maximum tolerated dose (MTD) of a drug is the highest dose of that drug that does not induce overt toxicity or undesirable side effects within a set time frame [63]. In the present study, all the compounds have low MTD being lower than 0.477 log (mg/kg/day) [35]. The oral rat chronic toxicity is the lowest dose of a drug that results in an observed adverse effect over a time period, while the oral rat acute toxicity or LD_50_ is the measurement of how much of a drug is required to kill 50% of rats in a test [35]. In terms of acute toxicity, Isopteropodin is the safest, while Strophanthidin is safest in terms of chronic toxicity. Similarly for toxicity to *Tetrahymena pyriformis*, Isopteropodin is the safest while for toxicity to Minnows, Strophanthidin is the safest. Despite the fact that the liver is the most common target organ for drug candidates in animal toxicity tests, hepatotoxicity seldom causes drug development to be halted during the preclinical stage. When a drug has great therapeutic promise, hepatotoxicity in humans may be tolerable due to the fact that it is frequently reversible and dose dependent [64]. In this study, only Strophanthidin is predicted to be non-hepatotoxic.

### Analyses of time-resolved trajectories

The most commonly used quantitative metric of similarity between two stacked protein structures (the reference and target structures) is the RMSD. It calculates the differences in distances between atoms in two overlaid structures, with a result of 0.0 indicating perfect overlap [65]. Over the 2-nanosecond trajectory, the BrMelMetRS-OOU and BrMelMetRS-Isopteropodin complexes showed marginally greater distortion than the BrMelMetRS-Strophanthidin complex in terms of variations in the RMSD of the Cα atomic coordinates. This is evidenced by the values of highest RMSD peak, the total RMSD, and the average RMSD. All the RMSD slopes induced by the holo forms show a gentle upward trend suggesting greater values with more simulation time. In this study, as it concerns RMSD peaks distribution patterns, all the holo forms show similar stability [66]. The structure and dynamics of proteins play a big role in how well they work. The motions of the residues are used to assess a protein’s mobility. The Root mean square fluctuation (RMSF) measures the structural flexibility of the protein by calculating the fluctuations of residues during molecular dynamics simulation [67,68]. While BrMelMetRS-Isopteropodin complex showed the greatest fluctuations amongst the holo structures at the global level, the BrMelMetRS-Strophanthidin complex showed the greatest fluctuations at the regional level (Pocket 1).

The PCA is used to statistically evaluate the various structural conformations of a protein generated during trajectories [69]. This study found that the BrucMetRS - Isopteropodin complex has the greatest global (total and average) motions of any holo structures, closely followed by the BrMelMetRS-OOU complex. At Pocket 1, the BrMelMetRS-OOU complex showed the highest regional (total and average) motions whereas, the BrMelMetRS-Strophanthidin complex showed greater regional motions than the BrucMetRS-Isopteropodin complex. Specifically, based on the highest motions, the best global and regional conformations are PC3, PC2, and PC2 for the BrMelMetRS-OOU, BrucMetRS–Isopteropodin, and the BrMelMetRS-Strophanthidin complexes respectively.

The B-factor is a measurement of a protein’s thermal stability based on the variation in atom locations in relation to average atomic coordinates [70]. Of all the holo structures, the BrMelMetRS-OOU complex showed the highest B-factor suggesting the greatest thermal instability. However, BrMelMetRS-Strophanthidin complex showed greater thermal instability than the BrucMetRS - Isopteropodin complex as seen by the global and regional average B factor values. The radius of gyration is the determinant of the compactness of the apo or holo protein during molecular dynamics simulation. It is a measure of the mass of atoms in relations with the protein complex centre of mass [71]. In terms of RoG along the trajectory, Isopteropodin induced the least compactness on the target (Figure 11, Table 6)

The dynamic cross-correlation map depicts the atomic correlation pattern in protein dynamics [72]. Of the 31 residues of the Pocket 1, the BrMelMetRS-OOU complex showed the highest number of anti-correlating residues. The BrMelMetRS-Strophanthidin and the BrucMetRS - Isopteropodin complexes have the same number of anti-correlating residues. The net values for all the residues in the Pocket 1 reveal that the Strophanthidin had the greatest anti-correlation effect on the target protein suggesting the greatest inhibitory activity at that site [46].

### BLAST

Many drugs are quite promiscuous and they would bind to several targets with structural similarity [73]. Fortunately, the bacterial methionyl-tRNA synthetase (MetRS) enzyme, which is required for protein synthesis, differs significantly from the human cytoplasmic equivalent (HCE) and therefore the HCE would not be inhibited by the lead compounds [74]. However, there is a possibility that the lead compounds interact with the unnamed protein product, CBX51367.1 which though has less than 30% identity, but has an E value less than 10^−6^ sharing significant similarity with BrucMetRS [75].

Taken together, this study demonstrates the potential antibacterial effect of the reference compound OOU, and the leads compounds, Isopteropodin and Strophanthidin. However, OOU is slightly unsaturated therefore showing poor drug likeness and the ability to inhibit P-glycoprotein I and II proteins. Both Isopteropodin and Strophanthidin have shown acceptable pharmacokinetic properties with Isopteropodin showing superior oral absorbability. In terms of time-resolved trajectory of the apo and holo structures of the target, Strophanthidin induced the greatest molecular distortion at Pocket 1 as seen with the RMSF, PCA, B-factor and DCCM results.

Strophantidin is a cardiac glycoside found in the seed of edible plant, *Corchorus olitorius*,, and has been used in the treatment of congestive heart failure. It functions by inhibiting the membrane bound Na+/ K+ ATPase in the cardiac muscles [76,77]. This blockage leads to influx of calcium ions leading to an inotropic effect. This mechanism of action is dose-dependent (0.1 μmol/L and 0.5 μmol/L), as Strophanthidin can be potentially cardiotoxic through Ca2+ overload, diastolic dysfunction, and arrhythmias when administered above maximum dose [76]. The anticancer potential of Strophanthidin has also been identified as it inhibits the MAPK, PI3K/AKT/mTOR, and Wnt/β-Catenin Signalling Pathways [78]. Further experiments are required to ascertain whether sub-therapeutic doses of Strophanthidin can induce significant antibacterial effect *in vivo*.

Isopteropodin is an oxindole alkaloid isolated from the cat claw plant (*Uncaria tomentosa)* whose water-soluble extract significantly enhanced immune function by increasing Phytohemagglutinin (PHA) stimulated lymphocyte proliferation in splenocytes of rats [79,80]. The findings of this study suggest that the plants, *Corchorus olitorius*, and *Uncaria tomentosa* containing the lead compounds, Strophantidin and Isopteropodin respectively could be exploited to make antibiotics for the treatment of brucellosis.

## Conclusion

This study indicates that Isopteropodin and Strophanthidin have the capacity to block the *Brucella mellitensis* Methionyl-tRNA synthetase at Pocket 1. Therefore, they could treat human brucellosis and hence have a high potential for clinical development. *In vivo and in vitro* studies are required to determine the effectiveness and toxicity of the lead compounds.

## References

1. Baddour MM. Diagnosis of brucellosis in humans: a review. J Vet. Adv. 2012; 2(4): 149–156

2. Sarker MAS, Sarker RR, Begum MM, Shafy NM, Islam MT, Ehsan MA et al.,). Seroprevalence and Molecular Diagnosis of Brucella Abortus and Brucella Melitensis in Bangladesh. Bangladesh J Vet Med. 2016; 14(2): 221–226.

3. Ducrotoy MJ, Bertu WJ, Ocholi RA, Gusi AM, Bryssinckx W, Welburn S, Moriyón. Brucellosis as an emerging threat in developing economies: lessons from Nigeria. PLoS Negl Trop Dis. 2014; 8(7): e3008. https://doi.org/10.1371/journal.pntd.0003008

4. Corbel MJ. Brucellosis in humans and animals. 2006, http://www.who.int/csr/resources/publications/Brucellosis.pdf 2014. 5. 004.

5. Aworh MK, Okolocha E, Kwaga J, Fasina F, Lazarus D, Suleman I, Poggensee G, Nguku P, Nsubuga P, Human brucellosis: seroprevalence and associated exposure factors among abattoir workers in Abuja, Nigeria - 2011. The Pan Afr Med J. 2013; 16: 103. https://doi.org/10.11604/pamj.2013.16.103.2143

6. Chahota R, Sharma M, Katochl RC, Verma S, Singh MM, Kapoor V, Asrani RK. (). Brucellosis outbreak in an organized dairy farm involving cows and in contact human beings, in Himachal Pradesh, India. Vet Arh. 2003; 73(2): 95–102.

7. Sofian M, Aghakhani A, Velayati AA, Banifazl M, Eslamifar A, Ramezani A. Risk factors for human brucellosis in Iran: a case-control study. Int J Infect Dis: IJID: official publication of the International Society for Infectious Diseases. 2008; 12(2): 157–161. https://doi.org/10.1016/j.ijid.2007.04.019

8. Falade, S. A case of possible brucellosis relapse in a veterinarian. Trop Vet. 2002; 20(4): 226–230

9. Poulou A, Markou F, Xipolitos I, Skandalakis PN. A rare case of Brucella melitensis infection in an obstetrician during the delivery of a transplacentally infected infant. J Infect. 2006; 53(1): e39–e41. https://doi.org/10.1016/j.jinf.2005.09.004

10. Chenais E, Bagge E, Lambertz ST, Artursson K. Yersinia enterocolitica serotype O:9 cultured from Swedish sheep showing serologically false-positive reactions for Brucella melitensis. Infection ecology & epidemiology. 2012; 2: 10.3402/iee.v2i0.19027. https://doi.org/10.3402/iee.v2i0.19027

11. Corbel MJ. Brucellosis: an overview. Emerg Infect Dis. 1997; 3(2): 213–221. https://doi.org/10.3201/eid0302.970219

12. Pathak AD, Dubal ZB, Karunakaran M, Doijad SP, Raorane AV, Dhuri RB, Bale MA, Chakurkar EB, Kalorey DR, Kurkure NV, Barbuddhe SB. Apparent seroprevalence, isolation and identification of risk factors for brucellosis among dairy cattle in Goa, India. Comp Immunol Microbiol Infect Dis. 2016; 47: 1–6. https://doi.org/10.1016/j.cimid.2016.05.004

13. Anyaoha CO, Majesty-Alukagberie LO, Ugochukwu ICI., Nwanta JA, Anene BM, Oboegbulam SI. Seroprevalencia y factores de riesgo de la brucelosis en perros de los Estados Enugu y Anambra, Nigeria. Rev Med Vet. 2020; 1(40): 5.

14. Ghodasara S, Roy A, Rank DN, Bhanderi BB Identification of Brucella spp. from animals with reproductive disorders by polymerase chain reaction assay. Buffalo Bull. 2010; 29(2): 98–108.

15. Alumasa L, Thomasid LF, Amanya F, Njorogeid SM, Moriyónid I, Makhandiaid J, Rushtonid J, Fèvreid EM, Falzonid LC. Hospital-based evidence on cost-effectiveness of brucellosis diagnostic tests and treatment in Kenyan hospitals. PLoS Negl Trop Dis. 2021; 15(1): 1–19. https://doi.org/10.1371/journal.pntd.0008977

16. Adesokan HK, Alabi PI, Stack JA, Cadmus SIB. Knowledge and practices related to bovine brucellosis transmission amongst livestock workers in Yewa, south-western Nigeria. J S Afr Vet Assoc. 2013; 84(1): Art N 121, 5 pages. https://doi.org/10.4102/jsava.v84i1.121

17. Smits HL, Cutler SJ. Contributions of biotechnology to the control and prevention of brucellosis in Africa. African Journal of Biotechnology. 2004; 3(12): 631–636.

18. McDermott JJ, Arimi SM. Brucellosis in sub-Saharan Africa: epidemiology, control and impact. Vet Microbiol. 2002; 90(1-4): 111–134. https://doi.org/10.1016/s0378-1135(02)00249-3.

19. OIE. (2009). Impact of Brucellosis on the Livestock Economy and Public Health in Africa. 18th Conference of the OIE Regional Commission For Africa, Ndjamena, Chad, 22-26 February 2009. Recommendation No. 2, 2, 204–205. http://www.oie.int/doc/ged/D6217. 2014. 9. 003

20. Gross A, Bouaboula M, Casellas P, Liautard JP, Dornand J. Subversion and utilization of the host cell cyclic adenosine 5’-monophosphate/protein kinase A pathway by Brucella during macrophage infection. J Immunol (Baltimore, Md. : 1950). 2003; 170(11): 5607–5614. https://doi.org/10.4049/jimmunol.170.11.5607

21. Marianelli C, Ciuchini F, Tarantino M, Pasquali P, Adone R. Genetic bases of the rifampin resistance phenotype in Brucella spp. J Clin Microbiol. 2004; 42(12): 5439–5443. https://doi.org/10.1128/JCM.42.12.5439-5443.2004

22. Kumari M, Chandra S, Tiwari N, Subbarao N. High Throughput Virtual Screening to Identify Novel natural product Inhibitors for MethionyltRNA-Synthetase of Brucella melitensis. Bioinformation. 2017; 13(1): 8–16. https://doi.org/10.6026/97320630013008

23. Oloso NO, Fagbo S, Garbati M, Olonitola SO, Awosanya EJ, Aworh MK, Adamu H, Odetokun IA, Fasina FO. Antimicrobial Resistance in Food Animals and the Environment in Nigeria: A Review. Int J Environ Res Public Health. 2018; 15(6): 1284. https://doi.org/10.3390/ijerph15061284

24. Bayram Y, Korkoca H, Aypak C, Parlak M, Cikman A, Kilic S, Berktas M. Antimicrobial susceptibilities of Brucella isolates from various clinical specimens. Int J Med Sci. 2011; 8(3): 198–202. https://doi.org/10.7150/ijms.8.198

25. Deniziak MA, Barciszewski J. Methionyl-tRNA synthetase. Acta Biochim Pol. 2001; 48(2): 337–350.

26. Ojo KK, Ranade RM, Zhang Z, Dranow DM, Myers JB, Choi R, Nakazawa HS, Edwards TE, Davies DR, Lorimer D, Boyle SM, Barrett LK, Buckner FS, Fan E, Van Voorhis WC. Brucella melitensis Methionyl-tRNA-Synthetase (MetRS), a Potential Drug Target for Brucellosis. PloS One, 2016; 11(8): e0160350. https://doi.org/10.1371/journal.pone.0160350

27. Berman HM, Westbrook J, Feng Z, Gilliland G, Bhat TN, Weissig H, Shindyalov IN, Bourne PE. The Protein Data Bank. Nucleic Acids Res. 2000; 28(1): 235–242. https://doi.org/10.1093/nar/28.1.235

28. Waterhouse A, Bertoni M, Bienert S, Studer G, Tauriello G, Gumienny, R, Heer, FT, de Beer T, Rempfer C, Bordoli L, Lepore R, Schwede T. SWISS-MODEL: homology modelling of protein structures and complexes. Nucleic Acids Res. 2018; 46(W1): W296–W303. https://doi.org/10.1093/nar/gky427.

29. Schrödinger L, DeLano W. PyMOL. (2020). Retrieved May 10, 2021 from http://www.pymol.org/pymol

30. Willard L, Ranjan A, Zhang H, Monzavi H, Boyko RF, Sykes BD, Wishart DS. VADAR: a web server for quantitative evaluation of protein structure quality. Nucleic Acids Res. 2003; 31(13): 3316–3319. https://doi.org/10.1093/nar/gkg565

31. Chen VB, Arendall WB, 3rd, Headd JJ, Keedy DA, Immormino RM, Kapral GJ, Murray LW, Richardson JS, Richardson DC. MolProbity: all-atom structure validation for macromolecular crystallography. Acta Crystallogr. Sect D, Biol Crystallogr. 2010; 66(Pt 1), 12–21. https://doi.org/10.1107/S0907444909042073

32. Kim S, Thiessen PA, Bolton EE, Chen J, Fu G, Gindulyte A, Han L, He J, He S, Shoemaker BA, Wang J, Yu B, Zhang J, Bryant SH. PubChem Substance and Compound databases. Nucleic Acids Res. 2016; 44(D1): D1202–D1213. https://doi.org/10.1093/nar/gkv951

33. Dallakyan S, Olson AJ. Small-molecule library screening by docking with PyRx. Methods Mol Biol (Clifton, N.J.), 2015; 1263: 243–250. https://doi.org/10.1007/978-1-4939-2269-7_19

34. Daina A, Michielin O, Zoete V. SwissADME: a free web tool to evaluate pharmacokinetics, drug-likeness and medicinal chemistry friendliness of small molecules. Scientific Rep. 2017; 7: 42717. https://doi.org/10.1038/srep42717.

35. Pires DE, Blundell TL, Ascher DB. pkCSM: Predicting Small-Molecule Pharmacokinetic and Toxicity Properties Using Graph-Based Signatures. J Med Chem. 2015; 58(9): 4066–4072. https://doi.org/10.1021/acs.jmedchem.5b00104

36. Molinspiration. Calculation of Molecular Properties and Bioactivity Score. 2015: Available at http://www.molinspiration.com/cgi-bin/properties

37. Salentin S, Schreiber S, Haupt VJ, Adasme MF, Schroeder M. PLIP: fully automated protein-ligand interaction profiler. Nucleic Acids Res. 2015: 43(W1), W443–W447. https://doi.org/10.1093/nar/gkv315

38. Hospital A, Andrio P, Fenollosa C, Cicin-Sain D, Orozco M, Gelpí, JL. MDWeb and MDMoby: an integrated web-based platform for molecular dynamics simulations. Bioinformatics (Oxford, England). 2012; 28(9): 1278–1279. https://doi.org/10.1093/bioinformatics/bts139

39. Afgan E, Baker D, Batut B, van den Beek M, Bouvier D, Cech M, Chilton J, Clements D, Coraor N, Grüning BA, Guerler A, Hillman-Jackson J, Hiltemann S, Jalili V, Rasche H, Soranzo N, Goecks J, Taylor J, Nekrutenko A, Blankenberg D. The Galaxy platform for accessible, reproducible and collaborative biomedical analyses: 2018 update. Nucleic Acids Res. 2018; 46(W1): W537–W544. https://doi.org/10.1093/nar/gky379

40. UniProt: the universal protein knowledgebase in 2021. Nucleic Acids Res. 2021; 49:D1 (2021)

41. National Center for Biotechnology Information (NCBI)[Internet]. Bethesda (MD): National Library of Medicine (US), National Center for Biotechnology Information; [1988] – [cited 2021 Jul 06]. Available from: https://www.ncbi.nlm.nih.gov.

42. Burra PV, Zhang Y, Godzik A, Stec B. Global distribution of conformational states derived from redundant models in the PDB points to non-uniqueness of the protein structure. Proceedings of the National Academy of Sciences of the United States of America. 2009; 106(26): 10505–10510. https://doi.org/10.1073/pnas.0812152106

43. Benkert P, Biasini M, Schwede T. Toward the estimation of the absolute quality of individual protein structure models. Bioinformatics (Oxford, England). 2011; 27(3): 343–350. https://doi.org/10.1093/bioinformatics/btq662

44. Biasini M, Bienert, S, Waterhouse A, Arnold K, Studer G, Schmidt T, Kiefer F, Gallo CT, Bertoni M, Bordoli L, Schwede T. SWISS-MODEL: modelling protein tertiary and quaternary structure using evolutionary information. Nucleic acids research. 2014; 42(Web Server issue), W252–W258. https://doi.org/10.1093/nar/gku340

45. Athar M, Sona, A. N., Bekono, B. D., & Ntie-Kang, F.. 2. Fundamental physical and chemical concepts behind “drug-likeness” and “natural product-likeness”. In Fundamental Concepts. 2020; (pp. 55-80). De Gruyter.

46. Rowaiye AB, Olubiyi J, Bur D, Uzochukwu IC, Akpa A, Esimone CO. In Silico Screening and Molecular Dynamic Simulation Studies of Potential Small Molecule Immunomodulators of the KIR2DS2 Receptor. J Phytomedicine Ther. 2021a; 20(1): 542–567.

47. Lovering F, Bikker J, Humblet C. Escape from flatland: increasing saturation as an approach to improving clinical success. J Med Chem. 2009; 52(21): 6752–6756. https://doi.org/10.1021/jm901241e

48. Ritchie TJ, Ertl P, Lewis R. The graphical representation of ADME-related molecule properties for medicinal chemists. Drug Discov. 2011; 16(1-2): 65–72. https://doi.org/10.1016/j.drudis.2010.11.002

49. Filimonov DA, Rudik AV, Dmitriev AV, Poroikov VV. Computer-Aided Estimation of Biological Activity Profiles of Drug-Like Compounds Taking into Account Their Metabolism in Human Body. International J Mol Sci. 2020; 21(20): 7492. https://doi.org/10.3390/ijms21207492

50. Yan A, Wang Z, Cai Z. Prediction of human intestinal absorption by GA feature selection and support vector machine regression. Int J Mol Sci. 2008; 9(10): 1961–1976. https://doi.org/10.3390/ijms9101961

51. Cui Q, Lu S, Ni B, Zeng X, Tan Y, Chen YD, Zhao H. Improved Prediction of Aqueous Solubility of Novel Compounds by Going Deeper With Deep Learning. Front Oncol. 2020; 10, 121. https://doi.org/10.3389/fonc.2020.00121

52. Peng Y, Yadava P, Heikkinen AT, Parrott N, Railkar A. Applications of a 7-day Caco-2 cell model in drug discovery and development. European J Pharm Sci: official journal of the European Federation for Pharmaceutical Sciences. 2014; 56: 120–130. https://doi.org/10.1016/j.ejps.2014.02.008

53. Hou T, Wang J, Li Y. ADME evaluation in drug discovery. 8. The prediction of human intestinal absorption by a support vector machine. Journal Chem Inf Model. 2007; 47(6): 2408–2415. https://doi.org/10.1021/ci7002076

54. Supe S, Takudage P. Methods for evaluating penetration of drug into the skin: A review. Skin Res Tech. 2021; 27(3): 299–308.

55. Prachayasittikul V, Prachayasittikul V. P-glycoprotein transporter in drug development. EXCLI journal. 2016; 15, 113–118. https://doi.org/10.17179/excli2015-768

56. Finch A, Pillans, P. P-glycoprotein and its role in drug-drug interactions. Aust Prescr, 2014; 37(4): 137–139.

57. Smith DA, Beaumont K, Maurer TS, D. L. Volume of Distribution in Drug Design. J Med Chem. 2015; 58(15): 5691–5698. https://doi.org/10.1021/acs.jmedchem.5b00201

58. Pardridge WM. Drug transport across the blood-brain barrier. J Cereb Blood Flow Metab: official journal of the International Society of Cerebral Blood Flow and Metabolism. 2012; 32(11): 1959–1972. https://doi.org/10.1038/jcbfm.2012.126

59. Zanger UM, Schwab M. Cytochrome P450 enzymes in drug metabolism: regulation of gene expression, enzyme activities, and impact of genetic variation. Pharmacol Ther. 2013; 138(1): 103–141. https://doi.org/10.1016/j.pharmthera.2012.12.007.

60. Horde GW, Gupta V. Drug Clearance. Treasure Island (FL): StatPearls Publishing. 2020.

61. Van Ness KP, Kelly EJ. Organic Cation Transporter 2. General Principles in Comprehensive Toxicology (Third Edition), 2018

62. Babcock JJ, Li M. hERG channel function: beyond long QT. Acta Pharmacol Sin. 2013; 34(3): 329–335. https://doi.org/10.1038/aps.2013.6

63. Stampfer HG, Gabb GM, Dimmitt SB. Why maximum tolerated dose?. Br J Clin Pharmacol. 2019; 85(10): 2213–2217. https://doi.org/10.1111/bcp.14032

64. Ballet F. Hepatotoxicity in drug development: detection, significance and solutions. J Hepatol. 1997; 26 Suppl 2: 26–36. doi: 10.1016/s0168-8278(97)80494-1. PMID: 9204407.

65. Carugo, O. How root-mean-square distance (rmsd) values depend on the resolution of protein structures that are compared. Journal Appl Crystallogr. 2003; 36(1): 125–128.

66. Rowaiye AB, Onuh OA, Sunday RM, Abdulmalik ZD, Bur D, Emeter NW et al., Structure-Based Virtual Screening and Molecular Dynamic Simulation Studies of the Natural Inhibitors of SARS-CoV-2 Main Protease. J Ong Chem Res. 2020; 5(1): 20–31.

67. Musyoka TM, Kanzi AM, Lobb KA, Tastan BÖ. Structure Based Docking and Molecular Dynamic Studies of Plasmodial Cysteine Proteases against a South African Natural Compound and its Analogs. Sci Rep. 2016; 6: 23690. https://doi.org/10.1038/srep23690

68. Hassan M, Shahzadi S, Seo SY, Alashwal H, Zaki N, Moustafa AA. Molecular Docking and Dynamic Simulation of AZD3293 and Solanezumab Effects Against BACE1 to Treat Alzheimer’s Disease. Front Comput Neurosci. 2018; 12: 34. https://doi.org/10.3389/fncom.2018.00034

69. Rowaiye AB, Onuh OA, Asala TM, Ogu AC, Bur D, Nwankwo EJ, Orji UM, Ibrahim ZR, Hamza J, Ugorji AL. In Silico Identification of Potential Allosteric Inhibitors of the SARS-CoV-2 Helicase. Trop J Nat Prod Res. 2021b; 5(1):165–177. https://doi.org/10.26538/tjnpr/v5i1.22

70. Parthasarathy S, Murthy MR. Protein thermal stability: insights from atomic displacement parameters (B values). Protein Eng. 2000; 13(1): 9–13. https://doi.org/10.1093/protein/13.1.9

71. Tannera JJ. Empirical power laws for the radii of gyration of protein oligomers. Acta Crystallogr. Section D, Struct Biol. 2016; 72(Pt 10: 1119–1129. https://doi.org/10.1107/S2059798316013218

72. De Vivo M, Masetti M, Bottegoni G, Cavalli A. Role of Molecular Dynamics and Related Methods in Drug Discovery. J Med Chem. 2016; 59(9): 4035–4061. https://doi.org/10.1021/acs.jmedchem.5b01684

73. Zhou H, Gao M, Skolnick J. Comprehensive prediction of drug-protein interactions and side effects for the human proteome. Sci Rep. 2015; 5: 11090. https://doi.org/10.1038/srep11090

74. Faghih O, Zhang Z, Ranade RM, Gillespie JR, Creason SA, Huang W, Shibata S, Barros-Álvarez X, Verlinde C, Hol W, Fan E, Buckner FS. Development of Methionyl-tRNA Synthetase Inhibitors as Antibiotics for Gram-Positive Bacterial Infections. Antimicrob Agents Ch. 2017; 61(11), e00999–17. https://doi.org/10.1128/AAC.00999-17

75. Pearson WR. An introduction to sequence similarity (“homology”) searching. Current protocols in bioinformatics. 2013; Chapter 3: Unit3.1. https://doi.org/10.1002/0471250953.bi0301s42

76. Bolognesi R, Cucchini F, Javernaro A, Zeppellini R, Manca C, Visioli O. Effects of acute K-strophantidin administration on left ventricular relaxation and filling phase in coronary artery disease. The Am J Cardiol. 1992; 69(3): 169–172. https://doi.org/10.1016/0002-9149(92)91298-i

77. Nakamura T, Goda Y, Sakai S, Kondo K, Akiyama H, Toyoda M. Cardenolide glycosides from seeds of Corchorus olitorius. Phytochemistry. 1998; 49(7): 2097–2101. https://doi.org/10.1016/s0031-9422(98)00421-x

78. Reddy D, Ghosh P, Kumavath R. Strophanthidin Attenuates MAPK, PI3K/AKT/mTOR, and Wnt/β-Catenin Signaling Pathways in Human Cancers. Front Oncol. 2020; 9: 1469. https://doi.org/10.3389/fonc.2019.01469

79. Sheng Y, Bryngelsson C, Pero RW. Enhanced DNA repair, immune function and reduced toxicity of C-MED-100, a novel aqueous extract from Uncaria tomentosa. J Ethnopharmacol. 2000; 69(2): 115–126. https://doi.org/10.1016/s0378-8741(99)00070-7

80. Honório ICG, Bertoni BW, Pereira AMS. Uncaria tomentosa and Uncaria guianensis an agronomic history to be written. Ciênc Rural. 2016; 46: 1401–1410.

